# A Modular Bioinstructive Platform for Additive-Free, Topography-Driven Stem Cell Differentiation and Patterning

**DOI:** 10.1101/2025.07.11.664383

**Authors:** Fatmah I. Ghuloum, Leo A.H. Zeef, Lee A. Stevens, Marco A.N. Domingos, Susan J. Kimber, Mahetab H. Amer

**Affiliations:** Division of Cell Matrix Biology and Regenerative Medicine, School of Biological Sciences, Faculty of Biology, Medicine and Health, The University of Manchester, UK; Department of Biological Sciences, Faculty of Science, Kuwait University, Kuwait; Bioinformatics Core Facility, Faculty of Life Sciences, The University of Manchester, UK; Low Carbon Energy and Resources Technologies Research Group, Faculty of Engineering, University of Nottingham, UK; Department of Mechanical and Aerospace Engineering, School of Engineering, Faculty of Science and Engineering & Henry Royce Institute, The University of Manchester, UK

**Author notes:** Corresponding author*: Dr Mahetab H. Amer.

**Keywords:** Differentiation, Hedgehog signalling, Mesenchymal stromal cells, Microparticles, Osteogenesis

## Abstract

Recreating 3D bone formation *in vitro* without biochemical inducers remains a longstanding challenge in preclinical testing. We present a scalable, bioinstructive platform based on polylactic acid microparticles with controlled dimpled surface features that direct mesenchymal stem cell differentiation through endogenous topography-mediated mechanotransduction, establishing a mechanistically validated, additive-free platform. These 3D topographical cues drive cytoskeletal reorganisation and induce osteogenesis via canonical Hedgehog signalling. RNA-Seq revealed early significant upregulation of cytoskeletal components and osteochondral transcription factors, including runt-related transcription factor 2 (*RUNX2*) and SRY-box transcription factor 9 (*SOX9*), followed by activation of the insulin growth factor-II pathway and osteogenic commitment. To demonstrate translational potential, two-photon polymerisation lithography was employed to engineer precisely-patterned 3D topographies, inducing graded GLI1 expression without added soluble cues. This establishes a modular, versatile platform for stem cell engineering, offering a topography-driven, non-genetic analogue to mechanogenetics with broad utility for regenerative medicine and human-relevant development of bone models.

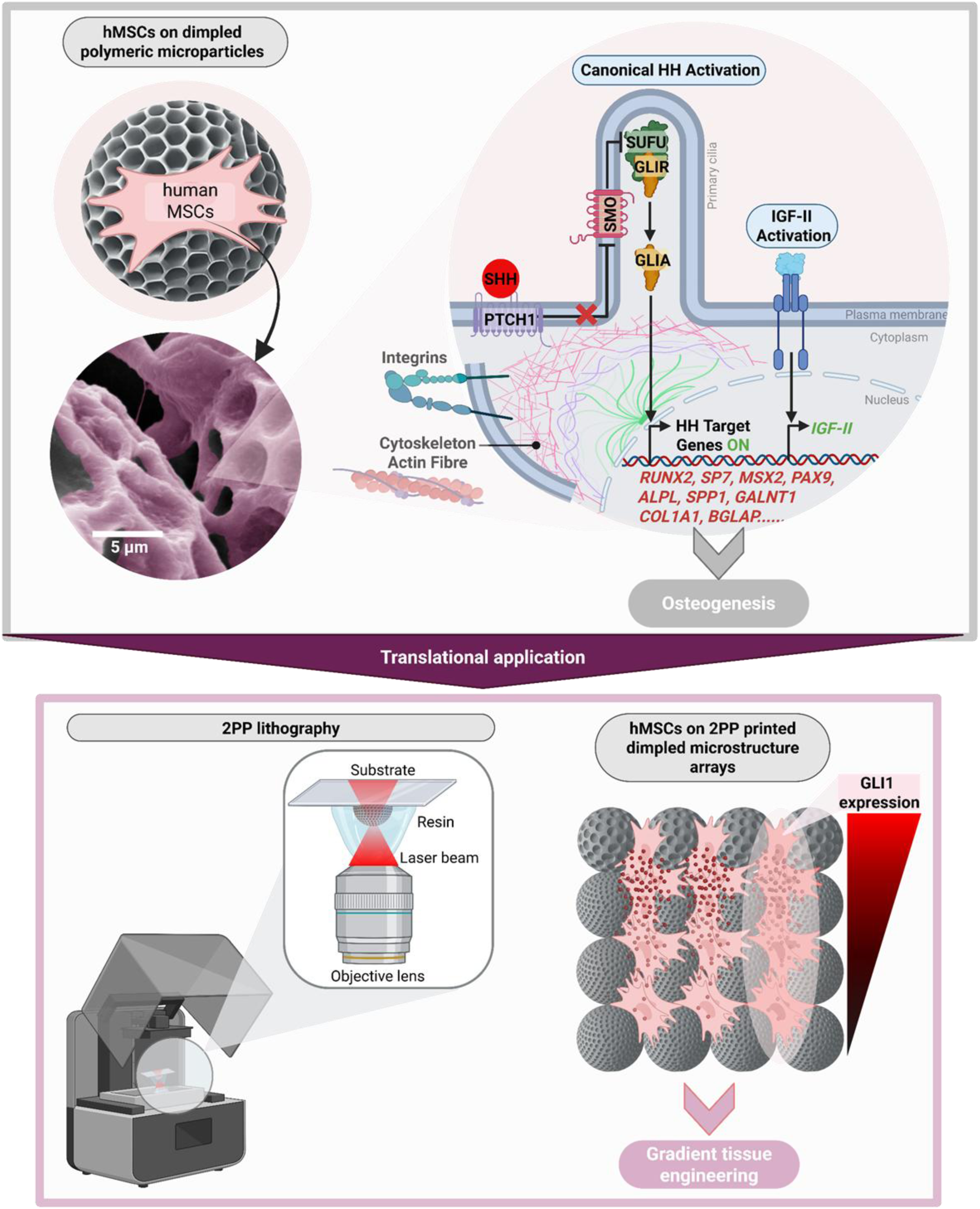
Graphical Abstract.

## Introduction

Bone formation is fundamental to skeletal development, homeostasis and repair, yet current tissue engineering strategies rely predominantly on costly growth factors that inadequately recreate the intricate processes of osteogenesis. Although it is well established that cell-matrix interactions can be used to direct cell response [1], a translation-ready approach to unlocking the potential of cell-instructive topographies for modulation of cell signalling has not been realised. This challenge is particularly relevant for tissue engineering, where traditional approaches rely heavily on costly, externally supplemented soluble factors that often fail to recapitulate the complexity of native tissue formation and can mask intrinsic cellular responses. The development of an alternative strategy leveraging cell-instructive biophysical cues to harness the inherent mechanosensory capabilities of cells to guide osteogenesis will transform regenerative medicine by enabling precise control over tissue formation while reducing costs.

Engineering functional bone tissue demands biomaterials that can precisely direct stem cell fate and patterning. While biochemical factors, such as dexamethasone, are widely used to induce differentiation of human mesenchymal stromal cells (hMSCs), they introduce confounding effects, which may result in inconsistent cellular responses and unintended lineage outcomes [2]. This includes potential activation of adipogenic pathways and upregulation of oxidative stress-related genes, compromising bone formation [3].

The hierarchical structure of bone’s extracellular matrix (ECM) has inspired scaffold design with customised topographies for bone regeneration [4]. Microtopographies, including pits, pillars, and gratings on titanium implants promote MSCs osteogenic differentiation in the presence of osteoinductive supplements [5, 6]. However, titanium’s stiffness and highly adhesive surface chemistry often favours fibrogenesis over the desired osteogenesis [7]. Similarly, tailoring topographically-textured micropatterns on glass slides coated with fibronectin (FN) has also been reported to promote osteogenic differentiation in rats [8], but FN suffers from low stability and degradation over seven days when subjected to mechanical forces such as laminar flow with shear stress [9], limiting long-term effectiveness in certain applications.

Recent advances in our understanding of mechanotransduction pathways have revealed promising alternatives to biochemical manipulation. The Hedgehog (HH) signalling pathway plays a pivotal role in the development of the skeletal system during embryogenesis [10], and regulates bone remodelling throughout postnatal life [11]. Critically, its mechano-responsiveness allows it to function as a central mediator of biomechanical cues, translating external mechanical forces into cellular response [12]. The core components of the HH signalling pathway consist of HH ligands, with Sonic Hedgehog (SHH) being the primary member, the twelve-pass transmembrane receptor, Patched1 (PTCH1), the seven-pass transmembrane signal transducer Smoothened (SMO), and the effector transcription factor Glioma-associated oncogene homolog 1 (GLI1) [13]. Dysregulation of the HH pathway has been reported in primary bone tumours such as osteosarcoma, and other musculoskeletal disorders such as osteoporosis and osteoarthritis [14], reinforcing its central role in skeletal homeostasis.

Topographically-textured 3D polymeric microparticles offer a promising platform for bone tissue engineering. mimicking native bone microarchitecture, while providing cell-instructive capabilities and a high surface area-to-volume ratio for large-scale cell expansion [15]. Previous research has demonstrated that 3D convex curvature enhances hMSC osteogenic differentiation by modulating the cytoskeleton, altering the distribution of focal adhesion proteins such as vinculin (VCL), and reducing stress fibre formation [16]. The osteoinductive impact of topographically-textured microparticles on murine mesenchymal progenitors is mediated by the SMO-dependent activation of GLI1, but this has not been assessed in relevant human primary cells [17].

This study aims to leverage these cell-instructive microparticles to investigate mechanically guided osteogenesis in primary hMSCs, mapping the transcriptional landscape of hMSCs cultured on these osteoinductive substrates to identify key bone-specific gene signatures and downstream molecular pathways governing lineage commitment and driving osteogenesis without the need for confounding biochemical supplements. Additionally, a key innovation of this study is the application of two-photon polymerisation (2PP) lithography to engineer precisely controlled GLI1 expression gradients, achieving fine-tuned modulation of cellular responses via engineered high-resolution topographies. While 2PP has been used to fabricate surfaces with gradients in roughness or stiffness [18, 19], its application to generate spatially defined gradients of cell-intrinsic signalling, such as GLI1 expression, represents a novel approach, bridging advanced fabrication techniques with the study of spatial regulation of cell fate. This study advances our understanding of how stem cells respond to 3D surface topography by showing that these surface-engineered microparticles with defined microfeatures can spatially modulate GLI1 expression in the absence of exogenous biochemical stimulation that may mask inherent cellular responses. Transcriptomic analysis further supports topography-induced activation of Hedgehog signalling and its role in directing early cell fate decisions. By leveraging high-resolution topographical cues to induce spatially patterned Hedgehog pathway activation, we present a versatile platform for probing the physical regulation of stem cell signalling and directing differentiation with microscale precision.

## Results

### Design and fabrication of surface-engineered, cell-instructive microparticles

To develop a scalable, supplement-free bone regeneration platform, we optimised our previously developed osteoinductive system [15, 17] for systematic investigation of topography-mediated mechanotransduction. This platform leverages topographical surface patterning to modulate cell responses. We engineered this platform based on three translational design criteria: (1) Microparticles should present precise microtopographical cues to enable spatially resolved, mechanotransduction-driven cell fate decisions; (2) Surface architecture should be tailored to promote optimal cell attachment and additive-free osteogenic differentiation [15, 17], in line with efforts to eliminate the need for exogenous biochemical cues for reduced regulatory complexity; and (3) Injectable particle size, facilitating minimally-invasive delivery and scalability for clinical translation.

Smooth and topographically-textured (‘dimpled’) microparticles were fabricated using a solvent evaporation oil-in-water emulsion technique, incorporating phase separation of fusidic acid (FA) as a sacrificial component to achieve defined surface patterning [15]. During hardening, FA is excluded from the bulk of the microparticle, producing distinctive surface patterns (Figure 1A). PLA was selected for its biocompatibility and hydrophobicity, preventing degradation during culture [15, 20].

**Figure 1:**
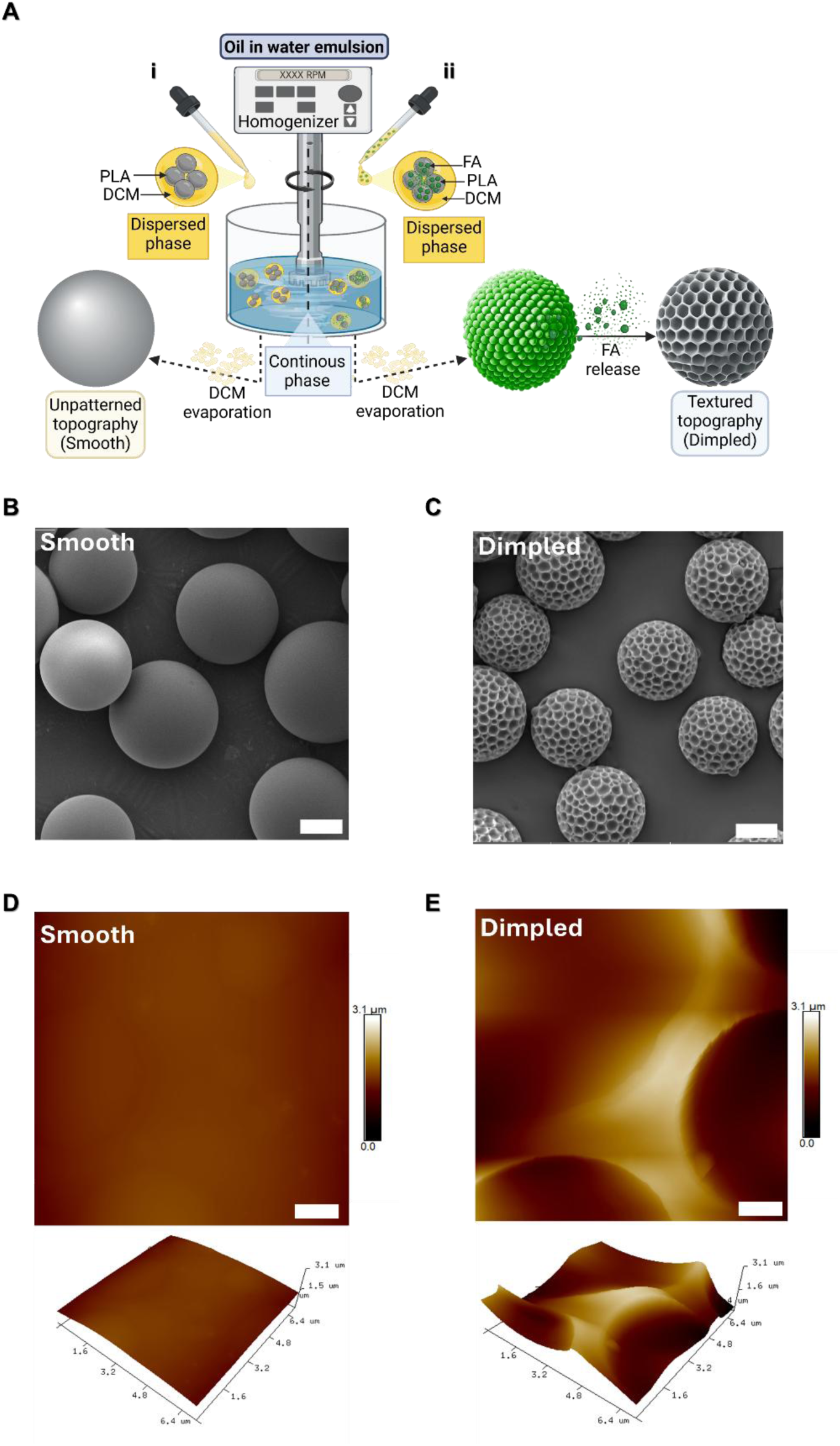
Fabrication of smooth and dimpled PLA microparticles. A) Schematic representation showing the fabrication workflow of smooth (i) and dimpled microparticles (ii) by a modified oil-in-water solvent evaporation emulsion method. B, C) Representative scanning electron microscopy images of smooth B) and dimpled C) microparticles acquired at 10 kV (Scale bars: 20 μm). D, E) Atomic force microscopy topography images of the smooth D) and dimpled E) microparticles (Scale bars: 1 μm). *Abbreviations: PLA, Poly(D,L-lactic acid); DCM, Dichloromethane; FA, Fusidic acid*.

Fabrication parameters were optimised to ensure comparable average sizes of the fabricated microparticles, providing smooth microparticles as an appropriate control for 3D surface topography. Distinct surface morphologies were achieved through modulation of processing parameters, including homogenisation speed, polymer concentration and polymer/FA ratio (Table 1).

**Table 1:**
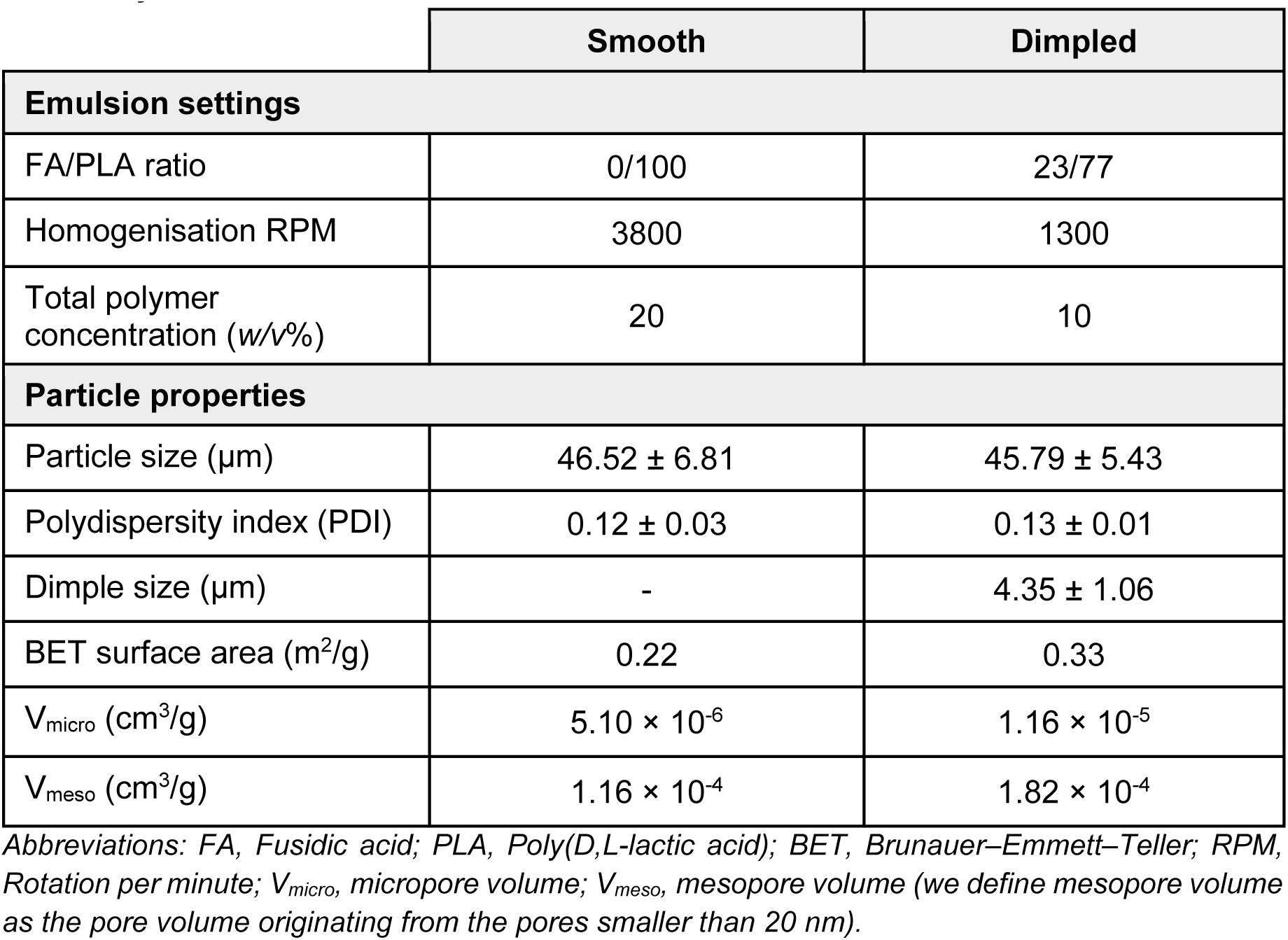
Fabrication parameters and properties of the polymeric microparticles used in this study.

Microparticles of similar size ranges of 46.52 ± 6.81 μm for smooth and 45.79 ± 5.43 μm for dimpled have been demonstrated to be injectable through clinically-relevant 21G needles [21]. The average dimple size falls within the mean size range that we previously reported to induce osteogenesis in hMSCs [15] and C3H10T1/2 cells [17] (Table 1). Surface area measurements (m²/g) indicated minimal porosity in both designs, reflected by low micro-pore (V_micro_) and meso-pore (V_meso_) volumes (Table 1). The successful fabrication of topographically-textured microparticles was confirmed through scanning electron microscopy (SEM) and atomic force microscopy (AFM) analyses, revealing well-defined surface patterns and feature dimensions (Figure 1). Dynamic light scattering measurements demonstrated narrow size distributions (Table 1). To maintain consistency in surface area available for cell attachment across samples, microparticle quantities were calculated to ensure uniform cell seeding density across 2D and 3D cultures.

Polymer microparticles are valuable for biomanufacturing workflows that depend on effective cell adhesion [15, 22]. hMSCs cultured on microparticles demonstrated excellent cell attachment and viability in serum-reduced medium (Figure 2A), with no significant differences observed between the two designs at any time point (Figure 2B). However, cell numbers in 3D-cultured samples were significantly lower than 2D-cultured controls at day 14 (*p* ≤ 0.0001).

**Figure 2:**
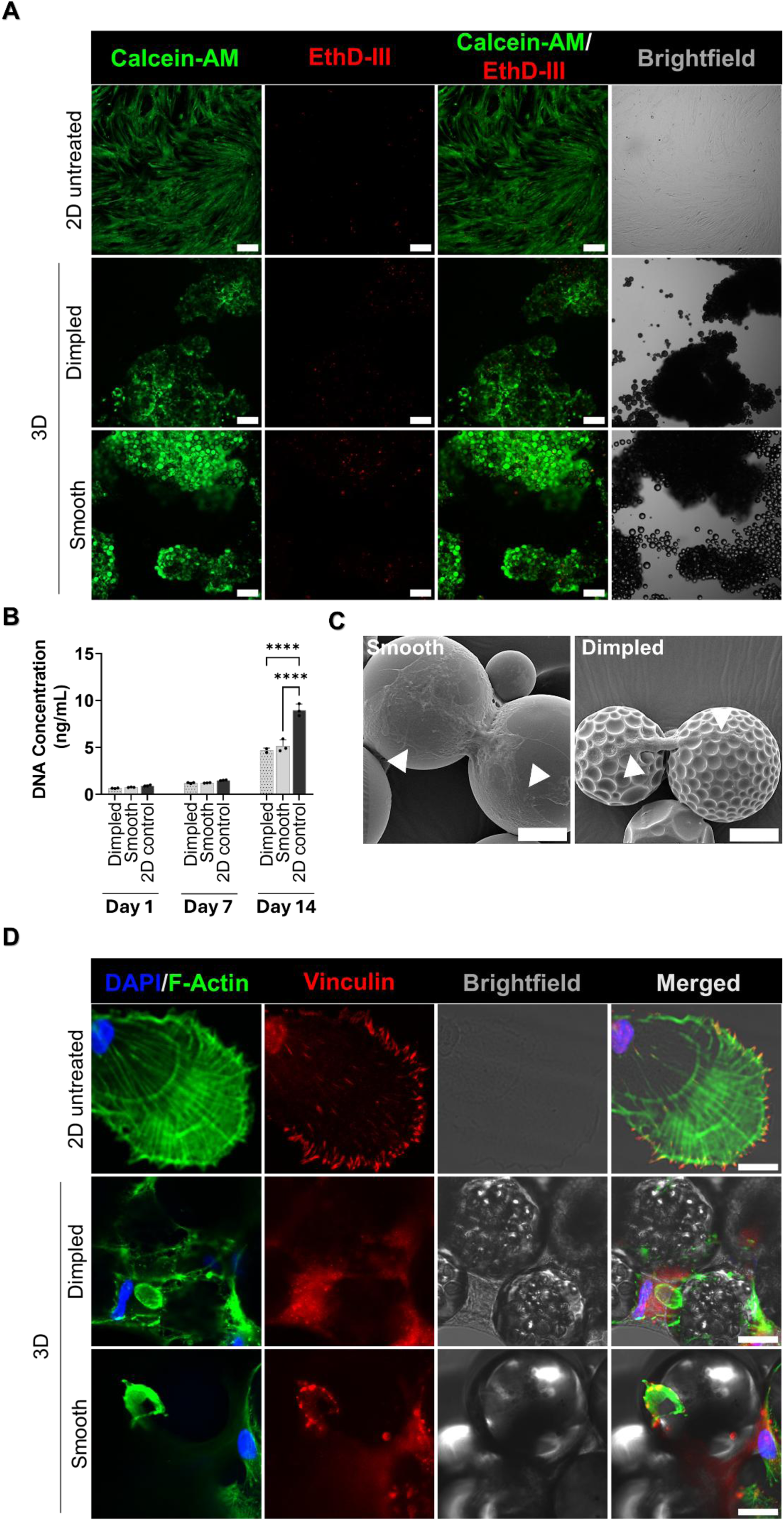
Impact of microparticles design on viability, proliferation, morphology and cytoskeletal organisation in hMSCs cultured on smooth and dimpled microparticles. A) Representative fluorescence microscopy images showing high viability of hMSCs on smooth and dimpled microparticles 3 days after seeding in serum-reduced medium. Live cells are stained with calcein-AM (green) and dead cells with ethidium homodimer III (EthD-III) (red) (Scale bars: 200 μm). B) DNA content in hMSCs cultured on dimpled and smooth microparticles versus 2D-cultured controls quantified at days 1, 7 and 14 days after seeding. Statistical significance is determined by two-way ANOVA with Tukey’s multiple comparisons test. Data represents mean ± SD (\*\*\*\**p* < 0.0001, N= 3 donors). C) Scanning electron microscopy images showing hMSCs morphologies 3 days after seeding on smooth and dimpled microparticles (Scale bars: 20 μm). D) Representative confocal maximum intensity projection images of hMSCs stained for VCL (red), F-actin (green), and nuclei (DAPI; blue) after 3 days in culture (Scale bars: 20 μm; N= 2 donors). White arrowheads indicate VCL localisation. *Abbreviations: EthD III, Ethidium homodimer III; VCL, Vinculin; F-actin, Filamentous actin; DAPI, 4′,6-Diamidino-2-phenylindole*

Cells demonstrated distinct morphological adaptations in response to topographical design, with a spread-out, flattened morphology on smooth microparticles and elongated, spindle-like appearance on the dimpled design (Figure 2C). This prompted the investigation of cytoskeletal architecture. Dimpled surfaces induced unique cytoskeletal arrangements (Figure 2D) that suggest differential mechanotransduction signalling. Apparent focal adhesions, which reflect cell-extracellular matrix interactions and subsequent cytoskeletal reorganisation [23], were examined by co-staining for VCL with F-actin. Immunostaining revealed well-defined, streak-like focal adhesions at the leading edge and pronounced F-actin stress fibres in hMSCs cultured on planar 2D-cultured controls. In contrast, cells seeded on dimpled microparticles exhibited diffuse VCL localisation and poorly defined focal adhesion structures, indicating impaired focal adhesion assembly [24]. Cells on smooth microparticles displayed mature focal adhesions (Figure 2D), highlighting the distinct effects of surface topography on focal adhesion dynamics.

### Transcriptome profiling identifies unique temporal transcriptional signatures of hMSCs cultured on dimpled 3D topographical features

While our previous work established the osteoinductive capacity of topographically-textured microparticles [15, 17] and the involvement of canonical HH signalling in this response in murine C3H10T1/2 cells, the underlying molecular mechanisms in human cells remained unexplored. Decoding how stem cells interpret osteoinductive physical cues at the transcriptional level will enable the design of bioinstructive systems that leverage mechanically-guided developmental mechanisms for regenerative and tissue engineering applications. To build on our findings, RNA sequencing (RNA-Seq) was therefore performed to determine the genome-wide transcriptional changes induced by different microparticle designs. hMSCs from three independent donors (Table S1) were seeded on smooth microparticles (serving as 3D topographical control) and two sets of dimpled microparticle cultures. On the following day, one set of dimpled samples was treated with KAAD-cyclopamine (a HH signalling inhibitor), generating three sample conditions: smooth microparticles, dimpled microparticles, and KAAD-cyclopamine-treated dimpled microparticles (Figure 3A).

**Figure 3:**
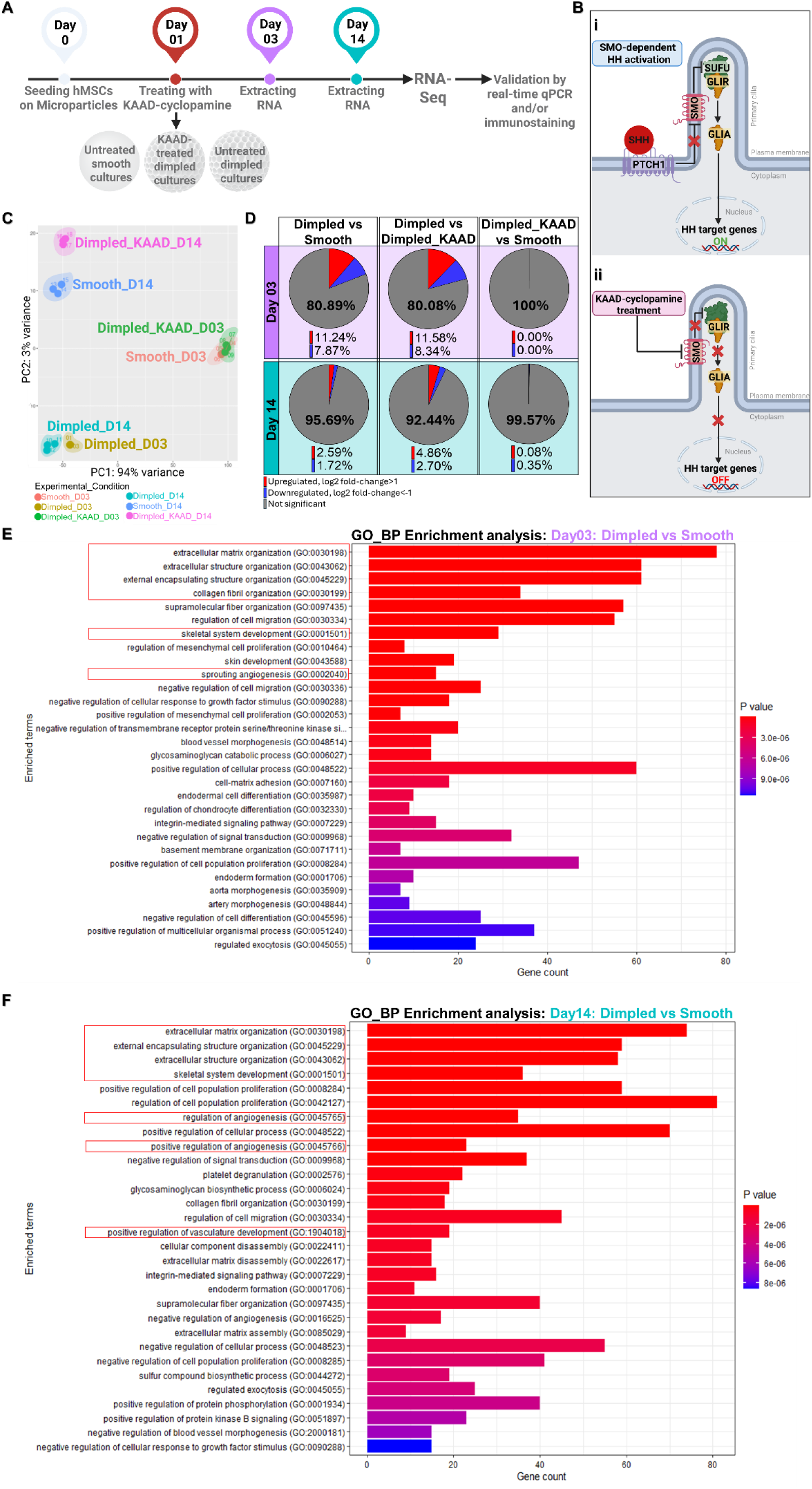
Dimpled microparticles show distinct transcriptional profiles compared to smooth micropartice cultures after 3 and 14 days in culture. A) Schematic representation of the experimental strategy for RNA-Seq utilised in this study. HMSCs from three donors were cultured on dimpled and smooth microparticles. On the following day one dimpled group was treated with KAAD-cyclopamine (HH antagonist). RNA-Seq was performed after 3 and 14 days in culture. B) (i) Schematic illustrating canonical Smoothened (SMO)-dependent HH activation, which begins when HH ligand inhibits Patched 1 (PTCH1), allowing SMO to overcome Suppressor of Fused (SUFU)-mediated repression and activates GLI, driving target gene transcription. (ii) KAAD-cyclopamine antagonises SMO, inactivating the canonical HH pathway. C) Principal component analysis plot displaying transcriptomic variance across dimpled, smooth and cyclopamine-treated dimpled samples at days 3 and 14 post-seeding. Each point represents an individual sample. D) Pie charts showing the average proportion of upregulated (log_2_ fold change > 1, red) and downregulated (log_2_ fold change < –1, blue) genes, with *p*_adj_ < 0.05, based on RNA-Seq analysis. E, F) Gene ontology enrichment analysis for biological processes in hMSCs cultured on dimpled versus smooth after 3 (E) and 14 (F) days in culture performed with Enrichr. Pathways of interest highlighted by red boxes. Length of the bar represents the degree of gene enrichment. Gene count is indicated on the x-axis. Colour indicates Benjamini-Hochberg adjusted *p* value. *Abbreviations: GLIR, Glioma-Associated oncogene homologs repressor form; GLIA, GLI activator form; PCA, Principal Component Analysis; GO, Gene Ontology; BP, Biological Process*.

Canonical HH signalling is initiated with the binding of SHH ligand to PTCH1, relieving the inhibition of the transmembrane protein SMO. The activation of SMO enables the nuclear translocation of the transcription factor GLI1, driving the transcription of HH target genes, such as *RUNX2* [25]. Treatment with KAAD-cyclopamine (SMO antagonist) inactivates this pathway (Figure 3B). RNA-Seq was performed at two time points: day 3, capturing any HH signalling activation and early topographically-induced osteogenic commitment, and at day 14, to assess HH signalling and downstream pathways governing cellular adaptation to the microenvironment (Figure 3A).

Principal component analysis (PCA) revealed distinct clustering by topographical condition and time point. At day 3, hMSCs cultured on dimpled microparticles segregated clearly from both smooth and cyclopamine-treated dimpled cultures (Figure 3C). The overlap observed between smooth and cyclopamine-treated dimpled samples suggests strong similarity of their transcriptional profiles at this early time point (Figure 3D), with only three genes being differentially expressed at day 3: Zinc finger protein 117 (*ZNF117*), poly(rc)-binding protein 1 (*PCBP1*), and thioredoxin domain containing 5 (*TXNDC5*). By day 14, cells cultured on dimpled microparticles formed distinct clusters relative to those on dimpled microparticles at day 3, indicating sustained transcriptional divergence driven by the dimpled topography. At day 14, only 0.43% of genes were differentially expressed between cells cultured on smooth microparticles and those on dimpled microparticle cultures treated with KAAD-cyclopamine (Figure 3D).

To identify putative drivers of hMSC adaptation to topography, we examined the top 30 up– and down-regulated DEGs at day 3 and day 14 post-seeding (Table S2 and S3, respectively). Early responses (day 3) were marked by upregulation of genes involved in cytoskeletal organisation, including tubulin alpha 3C (*TUBA3C*), and transcription factors linked to early osteoblastogenesis, such as GATA binding protein 4 (*GATA4*). By day 14, differentially expressed genes were increasingly associated with musculoskeletal tissue development and matrix remodelling, including sclerostin domain-containing 1 (*SOSTDC1*) and serpin family E member 2 (*SERPINE2*). Culture of primary hMSCs on dimpled microparticles resulted in 11979 genes (19.11%) at day 3 and 2701 genes (4.31%) at day 14 of all analysed genes (62700 genes) showing significant differential expression (with log_2_ fold change > 1 and *p*_adj_ < 0.05) relative to culture on unpatterned smooth microparticles.

Gene Ontology (GO) analysis was conducted to identify biological processes (BP) that were enriched in cells cultured on dimpled versus smooth microparticles. At day 3, enriched pathways were primarily related to ECM organisation, skeletal system development and angiogenesis (Figure 3E). By day 14, enrichment of ECM-related processes and skeletal system development remained prominent, with positive regulation of angiogenesis and vasculature development emerging (Figure 3F). Enriched GO terms with *p*_adj_-values and overlap metrics are listed in Table S4.

### Dimpled topographical features promote transcriptional upregulation of cytoskeletal genes and activate canonical Hedgehog signalling

At day 3 post-seeding, there was significant differential upregulation of key mechanosensing molecules, including piezo-type mechanosensitive ion channel component 2 (*PIEZO2*) and integrin subunit beta *4* (*ITGB4*), as well as associated structural genes such as *LAMA5*, which forms a functional complex with ITGB4 [26]. (Figure 4A).

**Figure 4:**
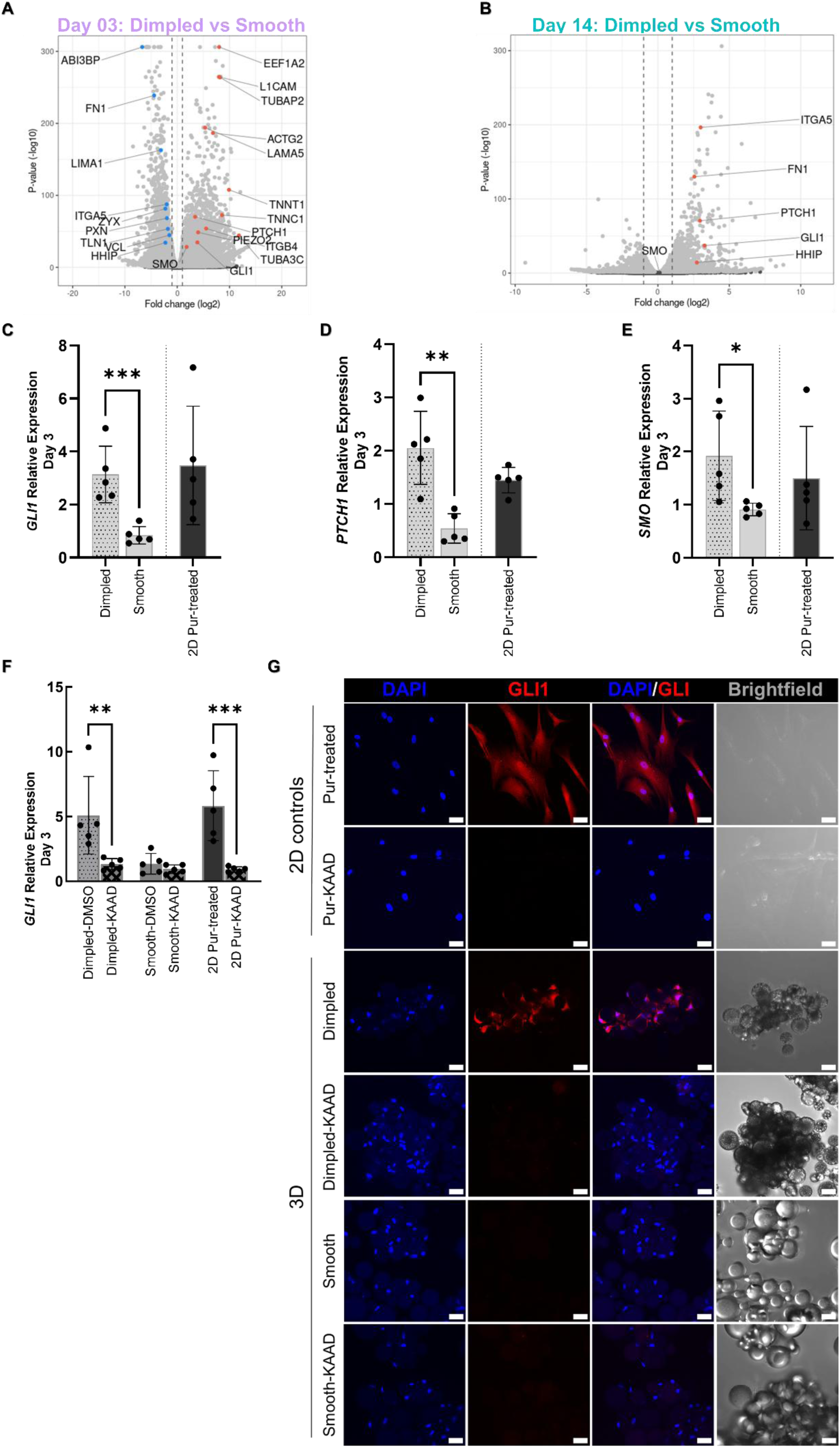
Culture of primary hMSCs on dimpled microparticles induces the upregulation of key Hedgehog signalling pathway components. A, B) Volcano plots displaying HH pathway-related differentially expressed genes in dimpled versus smooth microparticlecultures at day 3 (A) and day 14 post-seeding (B). Differential expression was defined as log*₂* fold change > 1 (upregulated, red) or < –1 (downregulated, blue) with *p*_adj_ < 0.05 (N = 3 donors)C-E) Quantitative real-time PCR (qPCR) analysis of key HH genes: *GLI1* (C), *PTCH1* (D), and *SMO* (E), after 3 days in serum-reduced medium, relative to untreated 2D controls. F) Relative *GLI1* expression after 3 days of 300 nM KAAD-cyclopamine treatment or 0.06% (*v/v*) DMSO in serum-reduced medium, and relative to 2D vehicle-only controls (with 0.06% DMSO). Expression in 2D purmorphamine-treated controls (2 μM) was calculated relative to 2D vehicle-only controls. (N= 5 donors). Statistical significance was calculated using one-way ANOVA with Tukey’s multiple comparisons test. Values are shown as mean ± SD (\**p* < 0.05, \*\**p* < 0.01, \*\*\**p* < 0.001). G) Representative confocal maximum intensity projection images of hMSCs in response to KAAD-cyclopamine treatment, stained for GLI1 (red) and nuclei in blue (DAPI) after 7 days in culture (N= 2 donors; Scale bars: 20 μm). Samples labelled with “-KAAD” indicate those treated with KAAD-cyclopamine. *Abbreviations: GLI1, glioma-associated oncogene homologs 1; SMO, smoothened; PTCH1, patched 1; HHIP, hedgehog-interacting protein; PIEZO2, piezo type mechanosensitive ion channel component 2; VCL, vinculin; TLN1, talin 1; PXN, paxillin; ZYX, zyxin; LAMIN1; LIM domain and actin binding 1; ACTG2, Actin Gamma 2, Smooth Muscle; TNNT1, troponin T1; TNNC1, troponin C1; EEF1A2, eukaryotic translation elongation factor 1 alpha 2; L1CAM, L1 cell adhesion molecule; TUBAP2, tubulin alpha pseudogene 2; TUBB4A, tubulin beta 4A; TUBA3C, tubulin alpha 3C; LAMA5, laminin subunit alpha 5; ABI3BP*, *ABI family member 3 binding protein; FN1, fibronectin 1; ITGA5, integrin subunit alpha 5; ITGB4, integrin subunit beta 4; DMSO, Dimethyl sulphoxide; DAPI, 4′,6-Diamidino-2-phenylindole; Pur, Purmorphamine; DIMP, Dimpled; SM, Smooth; DIMP_KD, Dimpled culture treated with KAAD-cyclopamine*.

The marked upregulation of eukaryotic translation elongation factor 1 alpha 2 (*EEF1A2*) and L1 cell adhesion molecule (*L1CAM*) suggests active cytoskeletal remodelling [27, 28]. Several tubulin genes were upregulated, including tubulin alpha pseudogene 2 (*TUBAP2*) and *TUBA3C*, which encode microtubule structural components. The upregulation of components of the actin–myosin contractile apparatus, including Troponin T1 (*TNNT1*) and actin gamma 2, smooth muscle *(ACTG2),* may reflect increased cytoskeletal tension in response to dimpled topography [29–31]. On the other hand, other ECM and cytoskeletal-associated genes, including *FN1,* ABI family member 3 binding protein (*ABI3BP*), and LIM domain and actin binding 1 (*LIMA1*) were downregulated. These elements play critical roles in regulating cytoskeletal dynamics by modulating focal adhesion and actin filament assembly in response to mechanical cues [32]. Consistent with this, focal adhesion-associated genes such as talin 1 (*TLN1*), paxillin (*PXN*) and *VCL* were also downregulated (Figure 4A). This aligns with our earlier data, where VCL immunostaining indicated apparent focal adhesion disassembly in hMSCs on dimpled microparticles (Figure 2D). Notably, *FN1* and integrin subunit alpha 5 (*ITGA5*) were significantly upregulated at day 14, suggesting temporal reorganisation of adhesion-related gene expression (Figure 4B).

Transcriptomic analysis at day 3 post-seeding revealed significant upregulation of key HH signalling genes such as *GLI1, SMO* and *PTCH1* (Figure 4A). The sustained activation of the HH signalling pathway was evident in dimpled microparticles after 14 days of culture (Figure 4B), although *SMO* expression was no longer differentially expressed relative to smooth microparticle cultures at day 14 (Figure 4B). Concurrently, the expression of hedgehog-interacting protein (*HHIP*), a negative regulator of HH signalling, was found to be significantly upregulated in dimpled microparticle-relative to smooth microparticle-cultures at day 14 post-seeding.

The differential expression of key HH pathway genes at day 3 was validated by real-time qPCR, confirming significant upregulation of *GLI1* (2.30-fold, *p* < 0.05; Figure 4C), *PTCH1* (1.51-fold; *p* < 0.0001, Figure 4D) and *SMO* (1.84-fold; *p* < 0.05, Figure 4E) in hMSCs on dimpled versus smooth microparticles, relative to 2D cultures.

To determine the route of HH signalling activation in dimpled microparticles-based cultures of hMSCs, GLI1 expression was analysed at the transcript and protein levels using real-time qPCR and immunostaining, respectively. At day 3 post-seeding, real-time qPCR revealed that the treatment of dimpled microparticle cultures with 300 nM KAAD-cyclopamine significantly reduced *GLI1* expression levels compared to untreated dimpled microparticle cultures (*p* < 0.01). This reduction was to a level comparable to that observed in hMSCs cultured on smooth microparticles (Figure 4F). This was conducted using a DMSO-containing vehicle control, which had no appreciable effects on GLI1 levels. Although multiple GO terms appear enriched in the comparison between KAAD-cyclopamine-treated dimpled and smooth surfaces, the overall differences are minimal, as indicated by the low gene counts per term and borderline significance (Figure S1). At day 7, immunostaining revealed that KAAD-cyclopamine effectively inhibited GLI1 upregulation at the protein level, while untreated dimpled cultures exhibited visibly higher GLI1 expression (Figure 4G). These findings highlight the dependence of HH pathway activation on SMO and confirmed the canonical activation of the HH signalling in hMSCs in response to 3D dimpled topographies.

Subsequent analyses focused on comparing dimpled against smooth microparticle cultures to isolate intrinsic effects of topographical features on hMSCs, as the cyclopamine-treated cultures closely resembled smooth microparticle cultures, and primarily served to confirm HH pathway involvement.

### Transcriptomic analysis identifies a transient, developmentally-relevant osteochondral state driving microparticle-induced osteogenesis

RNA-seq analysis revealed a distinct temporal progression in lineage commitment and differentiation of hMSCs, driven by the 3D dimpled topographies (Figure 5A).

**Figure 5:**
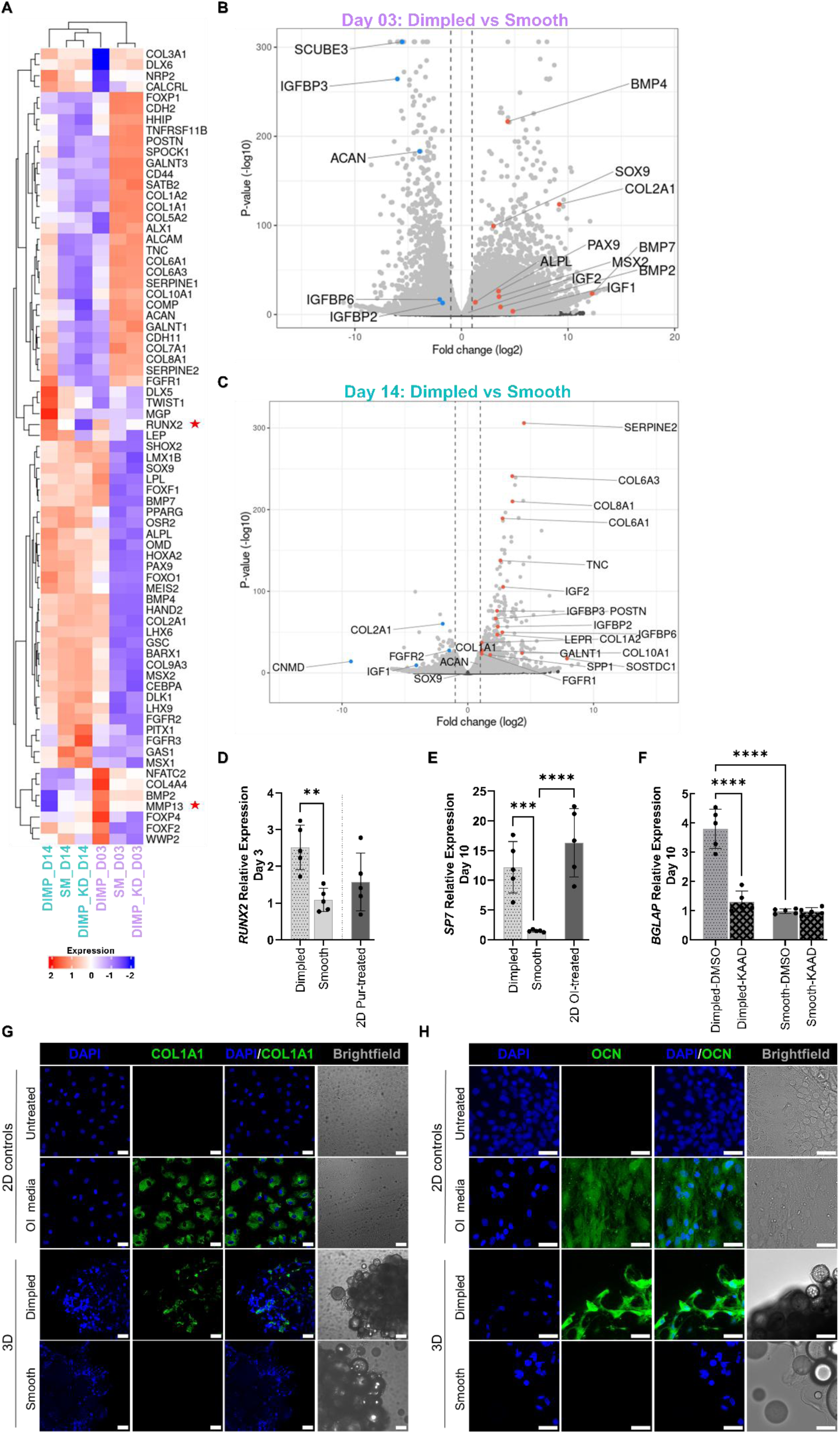
Transcriptomic analysis of hMSCs on dimpled versus smooth microparticles reveals novel insights into mechanically-guided osteogenesis without the confounding influence of biochemical additives. A) Heatmap showing expression of selected genes associated with skeletogenesis across experimental conditions at days 3 and 14 post-seeding. B, C) Volcano plots displaying key differentially expressed genes associated with cytoskeleton, osteogenesis, and chondrogenesis in dimpled versus smooth microparticle cultures at day 3 (B) and day 14 (C) post-seeding (log_2_ fold change > 1 and *p*_adj_ < 0.05, N= 3 donors). D, E) Relative quantitative real-time PCR (qPCR) analysis of gene expression at specific time-points in serum-reduced medium, relative to 2D controls: *RUNX2* (D; N= 5 donors) after 3 days, and *SP7* (E; N= 5 donors) after 10 days. 2D positive control treated with 2 μM purmorphamine is shown as a reference point for *RUNX2* expression. (F) Relative qPCR analysis of *BGLAP* expression after 10 days of either 300 nM KAAD-cyclopamine treatment or 0.06% (*v/v*) DMSO in serum-reduced medium, and relative to 2D vehicle-only controls (N= 5 donors). Statistical analysis was conducted using one-way ANOVA with Tukey’s multiple comparisons was applied. Values are presented as mean ± SD (\*\**p* < 0.01, \*\*\**p* < 0.001, \*\*\*\**p* < 0.0001). G, H) Representative maximum intensity projection confocal images of hMSCs stained for COL1A1 (green; G), and OCN (green; H) with nuclei counter-stained in blue (DAPI) after 10 days of culture (Scale bars: 50 μm). A single optical slice is used to represent brightfield for clarity. *Abbreviations: SOX9, SRY-box transcription factor 9; COL2A1, collagen type II alpha 1 chain; COL10A1, collagen type x alpha 1 chain; COL1A1, collagen type I alpha 1; COL6A1, collagen type VI alpha 1 chain; ACAN, aggrecan; RUNX2, runt-related transcription factor 2; SP7, specificity protein 7, also known as osterix; MSX2, Msh homeobox 2; SPP1, secreted phosphoprotein 1, also known as osteopontin; POSTN, periostin; PAX9, paired box 9; BGLAP, bone gamma-carboxyglutamate protein, also known as osteocalcin; ALPL, alkaline phosphatase; TNC, tenascin-c; SCUBE3, signal peptide-CUB-EGF domain-containing protein 3*; *SERPINE2, serine protease inhibitor 2; GALNT1, N-acetylgalactosaminyltransferase 1; LEPR, leptin receptor; SOSTDC1, sclerostin domain-containing protein 1; CNMD*, *chondromodulin; BMP, bone morphogenetic protein; IGF-II, insulin-like growth factor 2; IGFBP, insulin-like growth factor binding protein; DAPI, 4′,6-Diamidino-2-phenylindole; OI media, Osteoinductive media; Pur, Purmorphamine; DIMP, Dimpled; SM, Smooth; DIMP_KD, Dimpled cultures treated with KAAD-cyclopamine*.

The early mechanosensory response observed at day 3 post-seeding was accompanied by the significant upregulation of genes associated with early osteogenic commitment in hMSCs seeded on dimpled microparticles, including alkaline phosphatase (*ALPL*) and key transcriptional regulators of skeletogenesis such as Msh homeobox 2 (*MSX2*), and paired box 9 (*PAX9*). The expression of *RUNX2*, a key transcriptional regulator of osteogenesis, was observed in both smooth and dimpled microparticle cultures (Figure 5A). Several bone morphogenetic proteins (BMPs) that are critical to bone formation, including *BMP2*, *BMP4* and *BMP7*, were significantly upregulated in dimpled microparticle cultures (Figure 5B).

Interestingly, an upregulation of expression of SRY-box transcription factor 9 (*SOX9*), the master regulator of chondrogenesis, and its downstream target collagen type II alpha 1 chain (*COL2A1*) were also observed at day 3, suggesting a transient osteochondral bipotential state at this early time point [33] (Figure 5B). However, other chondrogenic-specific markers were also significantly downregulated at day 3, such as aggrecan (*ACAN*; Figure 5B) confirming incomplete commitment to chondrogenesis [34]. The substantial downregulation of signal peptide-CUB-EGF domain-containing protein 3 (*SCUBE3*) further suggests impaired BMP-mediated chondrogenesis [35] (Figure 5B).

By day 14, hMSCs on dimpled microparticles exhibited a clear trajectory towards a bone-specific transcriptional signature, displaying a profile indicative of osteogenic priming. This was supported by expression of *RUNX2* being sustained (Figure 5A) and significant upregulation of osteoblast-associated markers, including secreted phosphoprotein 1 (*SPP1*; encoding osteopontin), leptin receptor (*LEPR*), tenascin-C (*TNC*), *N-*acetylgalactosaminyltransferase *1* (*GALNT1*), and periostin (*POSTN*; osteoblast-specific factor 2). *SOSTDC1*, a HH-responsive BMP antagonist known to support trabecular bone maintenance [36], was also upregulated (Figure 5C). Upregulation of *FGFR1* but not *FGFR2* at this time point further supports osteogenic differentiation [37]. *SERPINE2* was among the most significantly upregulated genes (4.47-log_2_ fold change, *p*_adj_ < 0.01), consistent with its known role in extracellular matrix remodelling and osteogenic differentiation [38]. Moreover, bone matrix-associated collagens such as *COL1A1*, *COL6A1*, and *COL8A1* were significantly upregulated at day 14, suggesting commitment towards a skeletal or osteochondral lineage [39, 40]. Furthermore, differential upregulation of insulin-like growth factor 2 (*IGF-II*) along with its binding proteins *IGFBP2*, *IGFBP3* and *IGFBP6* was observed.

While significant upregulation of expression of *COL10A1* was observed at day 14, *SOX9* and *ACAN* were not differentially expressed at this time-point, while *COL2A1*, a cartilage-specific matrix protein [41] and chondromodulin (*CNMD*), a cartilage-associated glycoprotein [42], were significantly downregulated (Figure 5C). Another marker of hypertrophic chondrocytes, matrix metalloproteinase 13 (*MMP13*) [43], was detected at low levels at day 14; however, statistical significance could not be assessed due to limited read counts at this time point (Figure 5A).

Real-time qPCR validation demonstrated significant upregulation of expression of *RUNX2* (1.44-fold, *p* < 0.01; Figure 5D) at day 3 post-seeding and specificity protein *7* (*SP7*; 10.70-fold, *p* < 0.001; Figure 5E) at day 10 post-seeding in dimpled microparticle cultures compared to smooth cultures and relative to 2D controls. Notably, there was no significant difference in *SP7* expression between cells seeded on dimpled microparticles and in 2D cultures treated with osteoinductive medium. Moreover, elevated expression levels *SPP1* and integrin-binding sialoprotein (*IBSP*) were observed at day 10 post-seeding relative to 2D controls (Figure S2), though no statistically significant differences were detected. Immunostaining confirmed enhanced expression of COL1A1 (Figure 5G) and osteocalcin (OCN; Figure 5H) at day 10 post-seeding in dimpled versus smooth microparticle cultures.

To confirm the role of canonical HH signalling in topography-induced osteogenesis, the expression of bone gamma-carboxyglutamate protein (*BGLAP*), a late marker of osteogenesis, was examined after treatment with KAAD-cyclopamine. The expression *BGLAP*, encoding osteocalcin, was significantly reduced by 2.5-fold (*p* < 0.0001; Figure 5F) in dimpled cultures treated with KAAD-cyclopamine compared to untreated dimpled samples and relative to the 2D vehicle-only controls, confirming the dependence of its expression on canonical HH signalling.

### Dimpled topographical features induce temporal regulation of IGF-II expression

To elucidate downstream mechanisms of HH signalling and cross-talk governing the trajectory towards osteogenic commitment, QIAGEN’s Ingenuity^®^ Pathway Analysis (IPA) of DEGs in hMSCs cultured on different microparticle designs was performed to provide further insight into the dynamic molecular signatures associated with lineage commitment. IPA was performed on the set of 256 DEGs at day 14 post-seeding to identify the top canonical pathways predicted to be activated, offering insights into the regulatory networks orchestrating this osteogenic progression (Figure 6).

**Figure 6:**
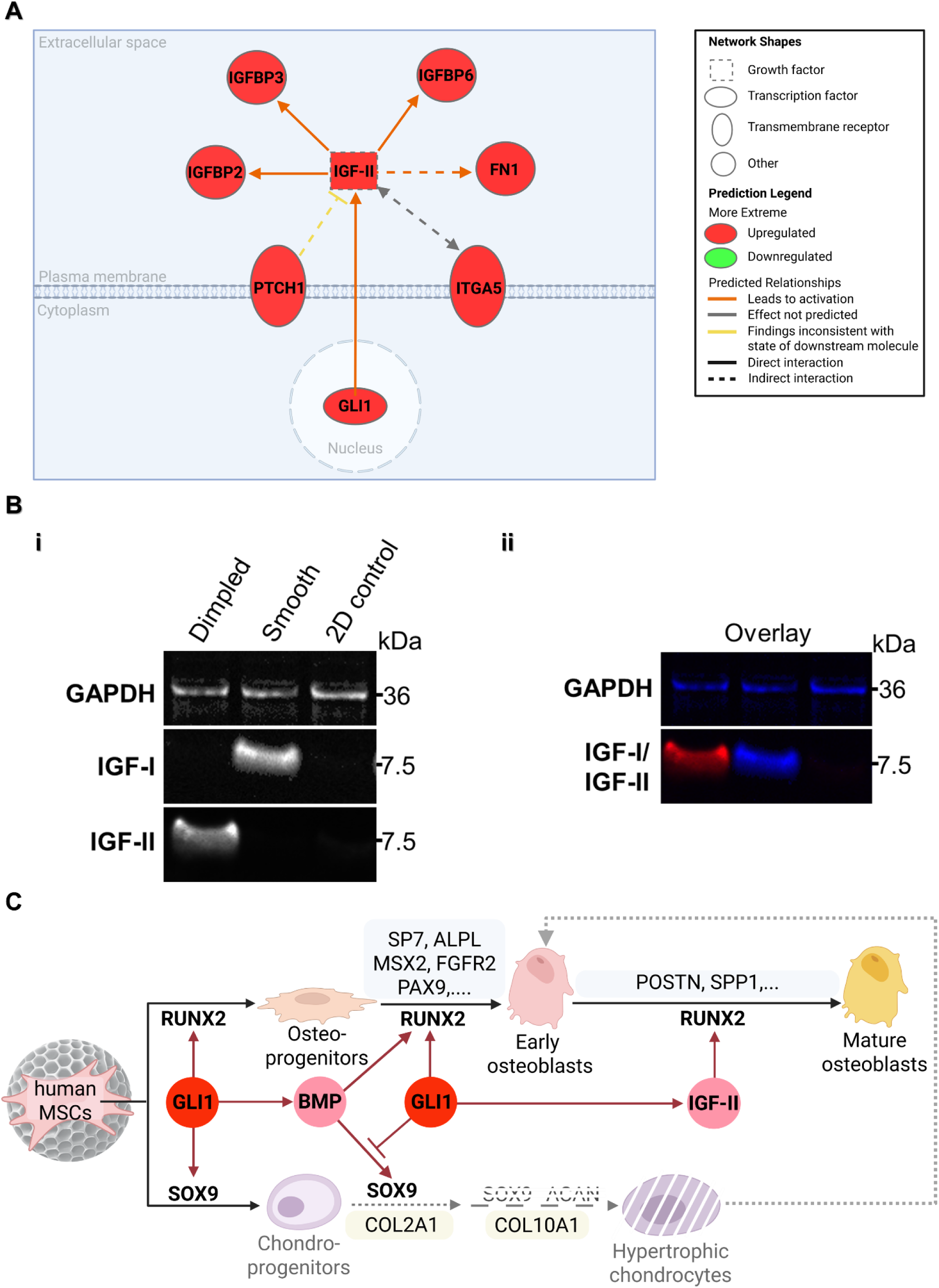
Dimpled microparticles promote expression of Insulin-like Growth Factor II (IGF-II) at day 14 post-seeding. A) Network analysis of the upstream regulatory network of insulin-like growth factor II (IGF-II) generated using IPA based on overlaid DEGs in hMSCs on dimpled versus smooth microparticles at day 14 post-seeding, highlighting transcriptional regulators, growth factors, and transmembrane receptors associated with IGF-II signalling. Nodes are colour-coded to represent expression levels: upregulated (red) and downregulated (green). Edges represent predicted relationships: activation (orange), findings inconsistent with expected relationships (yellow), and undefined effects (gray). This visualisation (created on BioRender.com) focuses specifically on upstream regulators of IGF-II, with full post-trimming analysis provided in supplementary data. B) Representative multiplexed fluorescence Western blot performed at day 14 post-seeding hMSCs on smooth and dimpled microparticles against 2D controls, showing individual channels (i), and the overlaid image (GAPDH and IGF-I in blue, IGF-II in red). (C) Schematic depicting the proposed mechanism of topographically-guided osteoinduction by 3D dimpled topographical features in hMSCs. *Abbreviations: IGFBP, insulin-like growth factor binding protein; FN1, fibronectin 1; PTCH1; Patched 1; GLI1, Glioma-associated oncogene homolog; ITGA5, Integrin subunit alpha 5; GAPDH, Glyceraldehyde-3-phosphate dehydrogenase*.

Canonical pathways were assessed using the activation z-score-(z-score > 2), which correlates observed gene expression with the expected direction of expression for DEGs [44]. IGF transport and uptake by IGFBPs was identified as the most significantly activated canonical pathway (z-score = 3.21, *p*_adj_ = 1.08 × 10^−11^, Table S5). Notably, *IGF-II* showed no differential expression at day 3 post-seeding (Figure 5B and Figure S3).

Interaction network analysis identified key regulatory molecules that may be responsible for the gene expression changes observed [45]. A direct activation link between *GLI1* and *IGF-II* expression was observed, while *PTCH1* indirectly inhibits *IGF-II* expression (Figure 6A). Despite *PTCH1*’s predicted negative influence, *IGF-II* remained activated, as indicated by the yellow node. Additionally, an indirect bidirectional interaction was identified between *IGF-II* and *ITGA5* (Figure 6A).

Experimental validation confirmed this striking dichotomy in IGF expression, where dimpled surfaces specifically induced IGF-II expression, while smooth surfaces promoted IGF-I expression (Figure 6B). The simultaneous expression of IGF-II and IGF-I proteins was confirmed using multiplex fluorescence-based Western blot analysis at day 14. This approach allowed concurrent detection of IGF-II and IGF-I, which possess a similar molecular weight, along with GAPDH as loading control, in the same sample [46]. This revealed the mature IGF proteins at their expected molecular weights, approximately 7.5 kDa [47] (Figure 6B). This topography-dependent regulation of IGF signalling suggests a novel mechanism by which 3D surface features direct cell fate. Dimpled topographies orchestrate the temporal progression of lineage commitment, initially establishing a bipotential osteochondral state driven by the activation of HH signalling, and transitioning toward osteogenesis via the IGF-II pathway (Figure 6C).

### Precision-engineered GLI1 expression gradients by two-photon lithography for spatially controlled stem cell signalling

Building on this mechanistic understanding of topography-induced signalling cascades, we fabricated precisely engineered micron-scale platforms. This proof-of-concept tested whether spatially-arranged microtopographies could generate defined zones of topographically-guided cellular signalling. This approach enables localised signalling control without genetic modification, serving as a novel physical analogue to mechanogenetics for controlling cell behaviour through material design.

2PP direct laser writing offers precision in fabricating intricate 3D microstructures while ensuring topographical consistency and uniformity [48]. Harnessing the osteoinductive capability of 3D dimpled topographical features, combined with the exceptional precision of 2PP, we engineered a GLI1 expression gradient by strategically employing cellular responses to topographical cues (Figure 7). GLI1 serves as a downstream transcriptional effector that reflects actual signal transduction and how cells translate topographical signals into differentiation responses [49].

**Figure 7:**
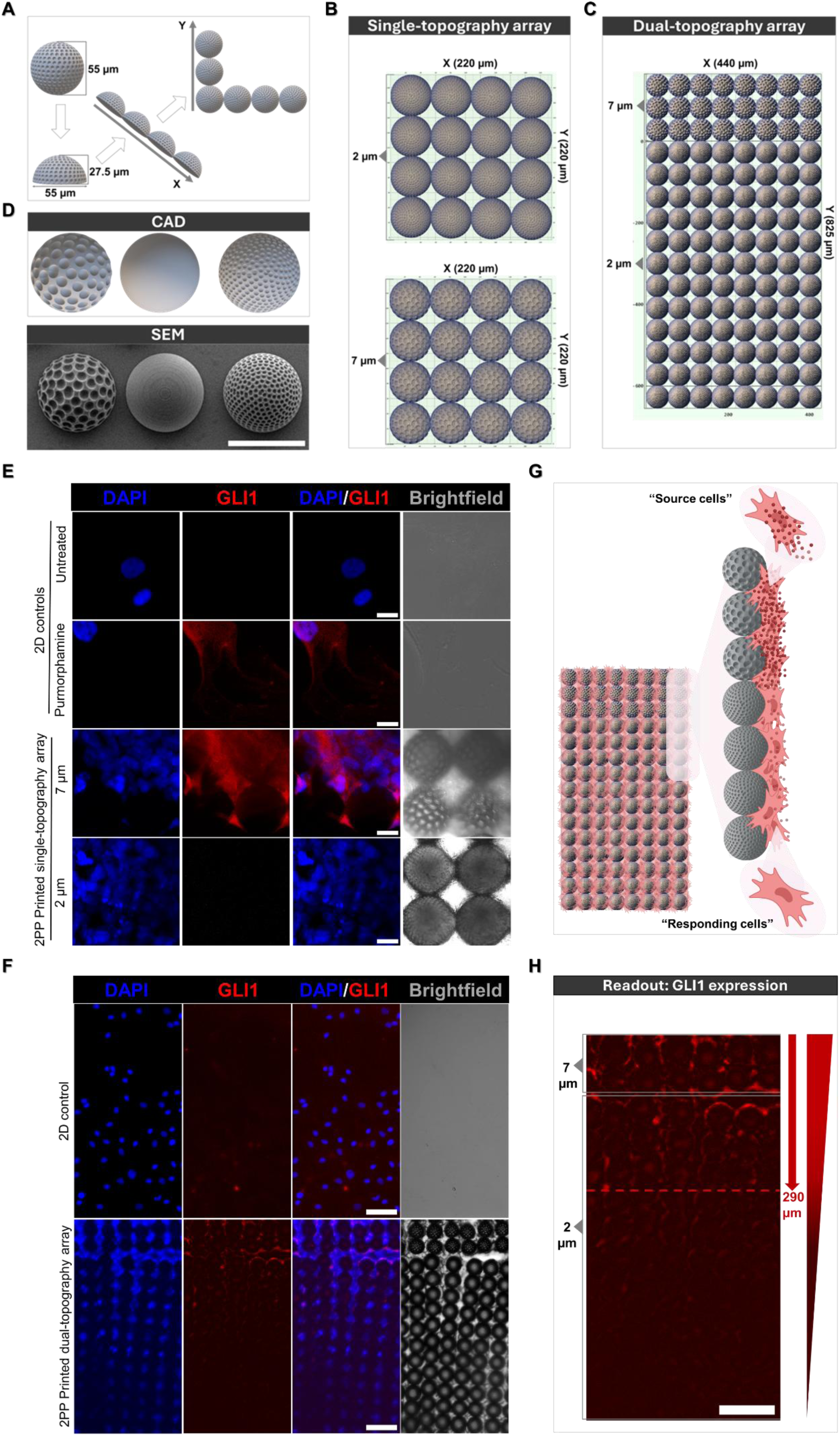
Engineering topography-induced GLI1 expression gradients using high-resolution two-photon polymerisation lithography. A) Schematic of the design strategy for 2PP fabrication. Hemispherical features (27.5 µm height and 55 µm width) served as building blocks that were systematically arranged in *x* and *y* directions to form structured arrays. B, C) Grid workflow language used to generate single-topography array (B) and dual-topography arrays combining 2 µm and 7 µm dimples (C). D) Computer-aided design of microparticles with different topographical features and corresponding SEM images after fabrication (Scale bar: 50 μm). E, F) Representative fluorescence images displaying GLI1 expression (red) and nuclei (DAPI; blue) in primary hMSCs seeded on single-topography arrays after 7 days in culture (E; scale bars: 20 µm; N= 2 arrays), and on dual-topography arrays featuring three rows of 7 µm dimpled hemispheres and twelve rows of 2 µm dimpled hemispheres (F, scale bar: 100 µm; N= 2 arrays). hMSCs cultured on glass slides served as 2D controls. DAPI channel is presented as maximum intensity projection images, and GLI1 as sum slice projection images. A single slice is used to represent brightfield for clarity. A sliding paraboloid background subtraction was applied to the red channel in dual-topography arrays, and standard background subtraction was applied for corresponding 2D controls. G) Schematic of the source-receiver configuration: MSCs seeded on 7 µm dimples act as ‘source cells’, while cells attaching to adjacent 2 µm dimpled rows are designated as ‘responding cells’ [Created with BioRender.com]. H) Representative confocal z-projection image (sum slices projection) displaying graded GLI1 expression, demonstrating the spatial correlation between topographical feature sizes and GLI1 activation (Scale bar: 100 µm). *Abbreviations: GLI1, Glioma-associated oncogene homolog 1; DAPI, 4′,6Diamidino-2-phenylindole*

Hemispherical microstructures (27.5 µm height) were fabricated with precisely controlled dimple sizes of either 2 µm or 7 µm (Figure 7A). These structures were imported into the DeScribe software (v2.7, Nanoscribe GmbH, Germany), and systematically arranged along the x and y axes to create two array configurations: x= 220 µm and y= 220 µm to achieve the single-topography array (Figure 7B), and x= 440 µm and y= 825 µm to engineer dual-topography arrays (Figure 7C). This approach minimised slicing-induced lines during 2PP fabrication, preventing artifacts. IP-Visio, a methacrylate-based commercial resin, was selected for its biocompatibility and lower autofluorescence compared to other photoresins [50]. Additionally, its reported stiffness (1.8 ± 0.64 GPa) [51] closely aligns with that of PLA microparticles [15], making it the optimal photoresin for this study. Precision of designs was validated by SEM imaging, demonstrating reproducible topographical features (Figure 7D). Initial studies using single-topography arrays revealed that 7 µm dimples resulted in a significant upregulation of GLI1 expression in hMSCs as previously observed with dimpled PLA microparticles, whereas 2 µm dimple sizes showed minimal effect (Figure 7E).

Inspired by this differential response, dual-topography arrays were then engineered. These arrays featured hemispherical structures with two dimple sizes: The top three rows featured 7 µm dimples, optimised to stimulate hMSCs as ‘source’ cells expected to induce GLI1 expression. In contrast, subsequent rows consisted of 2 µm dimpled structures, where hMSCs acted as ‘responding’ cells (Figure 7G). The platform was designed around the principle that physical microenvironmental features can orchestrate spatially regulated signalling responses, analogous to how patterned matrix cues guide tissue development [52]. Inspired by principles of localised activation and spatial propagation seen in developmental systems, we hypothesised that precisely engineered surface topographies could generate spatial heterogeneity in mechanotransduction responses. To test this, GLI1 expression in primary hMSCs seeded on this dual-topography array was evaluated at day 7 post-seeding. A visible spatial gradient of GLI1 expression emerged in the absence of any exogenous biochemical agonists (Figure 7F, H). Highest GLI1 expression was visibly highest within ∼290 µm of the ‘source’, gradually declining on more distant regions. This pattern persisted despite minor lateral displacements within the arrays during pre-seeding preparation, indicating robust spatial control. These findings demonstrate that engineered topographical features alone can induce spatially resolved intracellular signalling in stem cells, generating defined zones of mechanically-induced signalling. This provides a powerful platform for exploring spatial aspects of mechanotransduction and offers a biochemical-free approach for engineering zonally-patterned cellular responses, which is potentially transformative for developmental biology studies and regenerative material design.

## Discussion

Polymeric microparticles with engineered surface topographies offer powerful bottom-up engineering tools for directing stem cell fate through mechanical rather than biochemical cues [15, 20]. This study demonstrates that our topographically-engineered dimpled microparticles induce osteogenic differentiation in hMSCs by mechanical stimuli alone. Canonical HH signalling mediates this response, establishing a direct mechanotransductive mechanism. This materials-based approach offers a scalable, additive-free platform for developing cell-instructive 3D *in vitro* culture systems, with strong translational potential for regenerative medicine and stem cell biomanufacturing.

Topographical cues modulate mechanotransduction by altering cell morphology through cytoskeletal reorganisation [15, 24]. The elongated cell morphology observed on dimpled microparticles suggested increased cytoskeletal tension [53], in contrast to the isotropic spreading seen on smooth microparticles [54]. Elongated cell morphology is associated with altered focal adhesions, enhancing osteogenic potential in hMSCs [55]. Geometrical cues have been demonstrated to promote MSCs differentiation independent of soluble factors, with cytoskeletal-disrupting agents modulating these shape-based trends [56]. The increased expression of the mechanosensitive *ITGB4* at the earlier timepoint [57] is known to reduce VCL localisation within focal adhesions [58]. This is consistent with the significant downregulation of *VCL, PXN,* and *TLN1* and diffuse VCL distribution observed at day 3. Furthermore, the downregulation of LIM domain kinases, including *LIMA1* and *ZYX,* has been associated with actin depolymerisation by regulating actin filament turnover [59, 60]. The mechanosensitive ion channel *PIEZO2*, which is critical for proper SMO activation [61], was also significantly upregulated on dimpled topographies. Since substrate stiffness alone does not activate PIEZO2 [62], this suggests a topography-specific mechanical activation pathway. PIEZO2 activity supports calcium influx critical for SMO activation [63], potentially linking external mechanical cues to intracellular HH pathway responses. In our system, dimpled topographies coincided with disrupted actin organisation and diffuse VCL distribution, suggesting a mechanotransductive mechanism involving actin depolymerisation [17]. This is consistent with mechanisms reported during oestrogen withdrawal in murine osteocyte-like cells, where similar cytoskeletal changes were associated with HH activation [24].

Dimpled microparticles activated the canonical HH signalling pathway as early as day 3 post-seeding, with activation sustained through day 14. SMO-dependence of HH signalling activation in dimpled microparticle cultures was confirmed by the significantly reduced GLI1 expression in dimpled samples treated with KAAD-cyclopamine, and the similar transcriptional profile of cyclopamine-treated dimpled microparticle cultures to smooth microparticles. This aligns with our previous findings on murine embryonic mesenchymal progenitors [17]. While HH pathway components showed distinct regulation at day 3, gene expression profiles converged by day 14, suggesting that topography-driven signalling acts within a narrow temporal window. The significant downregulation of *BGLAP* in hMSCs cultured on dimpled microparticles and treated with KAAD-cyclopamine relative to untreated dimpled cultures confirmed the central role of HH signalling in topography-induced osteogenesis.

Significant upregulation of *HHIP* in dimpled cultures at day 14 suggests feedback attenuation of HH activity to maintain signalling homeostasis and prevent aberrant pathway activation [64]. As a pro-osteogenic gene, *HHIP* plays a pivotal role in regulating osteogenic mesenchyme in the coronal suture of mice and has been implicated in human embryonic skeletal development [65]. The lack of SMO differential expression at this stage is consistent with its known post-translational regulation, and could be attributed to the activation of IGF-I (Figure 6), which has been positively correlated with SMO expression in a GLI1-independent manner [66]. Moreover, sustained SMO expression results in impaired postnatal bone formation in mice [67]. This suggests that SMO activation is tightly regulated and transient during osteogenesis, in line with our findings.

The activation of the canonical HH pathway mirrors processes observed during skeletal development, where *GLI1*-expressing cells exhibit osteochondrogenic potential by contributing to endochondral and intramembranous ossification through the induction of *SOX9* and *RUNX2* expression, respectively [68]. It has been reported that the interaction of BMPs with the HH pathway shifts the balance towards *RUNX2* expression, driving MSCs commitment to osteoblasts [69]. BMP2 has also been suggested as a direct target of GLI1 [70], with GLI1 serving as a critical mediator between HH and downstream BMP pathway [71]. Early GLI1 expression induced both SOX9 and RUNX2, reflecting a transient bipotential osteochondroprogenitor state. While this early bipotential signature might suggest endochondral ossification, the absence of cartilage-specific matrix proteins, such as *ACAN,* despite transient *COL2A1* at day 3, argues against progression through a full endochondral ossification route. The significant upregulation of *COL10A1* at day 14, a marker of hypertrophic chondrocytes [43], likely reflects residual early chondrogenic activity rather than full hypertrophic transition. Further investigation is needed to clarify the role of *COL10A1* in this context. This initial bipotentiality is critical for the development of certain intramembranous bones via secondary cartilage [72]. Significant upregulation of *LEPR* on dimpled microparticles was observed at day 14, which aligns with reported *in vivo* findings that skeletal stem cells exhibit osteoblast-chondrocyte transitional identities in young bone marrow, progressively being replaced by *LEPR*-expressing stromal cells at later stages to serve as a source of osteoblasts in adult marrow [73].

*RUNX2* remained expressed across both microparticle designs at both time points, likely induced by matrix stiffness of the PLA microparticles [74]. This was accompanied by upregulation of osteogenic co-regulators *MSX2*, a transcription factor that synergises with *RUNX2* and drives *SP7* expression [75], and *PAX9*, a transcription factor required for craniofacial and tooth development and associated with early skeletogenesis, regulates key genes for bone formation such as *ALPL* and *COL1A1* [76]. These transcriptional regulators support osteoprogenitor proliferation, suppress alternative lineages, and promote expression of key drivers of osteoblast maturation.

MSCs differentiation to osteoblasts progressed with the subsequent expression of *SP7*, a downstream effector of *RUNX2* and master regulator of osteoblast differentiation [77], accompanied by increased expression of COL1A1 and OCN in hMSCs cultured on dimpled relative to smooth microparticles. By day 14 post-seeding, hMSCs demonstrated a clear trajectory towards osteogenesis. This was evidenced by the upregulation of *POSTN*, which is regarded as a marker of intramembranous ossification preceding increase in other osteogenic genes [78]. The expression of *TNC*, an ECM glycoprotein involved in osteogenesis and mineralisation [79], was also increased and is known to be induced by mechanical stimuli in murine pre-osteoblastic cell [80]. SERPINE2, essential for the early osteogenic commitment of MSCs [38], were also dramatically upregulated. Moreover, GALNT1 is critical for the expression of *SPP1* and *IBSP* in osteoblasts [81]. In addition, the transcriptional profile of hMSCs on dimpled microparticles revealed an ECM niche associated with early osteogenesis.

Collagen VI is a key pericellular matrix component found in MSCs, pericytes, and osteoprogenitors, supporting adhesion, survival, and mechanosensitivity [40]. COL8A1, more typically associated with vascular endothelium, was also upregulated. Its role in matrix remodelling and angiogenesis suggests a vascularised osteoprogenitor phenotype [82]. Gene ontology analysis confirmed enrichment in skeletal and vascular development pathways, reflecting the coupling of angiogenesis and intramembranous bone formation, where mesenchymal condensation centres facilitate blood vessel formation, enabling nearby mesenchymal cells to differentiate into osteoblasts [83]. The dominance of osteoblast-specific markers, validated on both genetic and protein levels, highlights the intramembranous ossification trajectory of hMSCs on dimpled microparticles. These results underscore the potential to integrate bone formation and vascular growth, paving the way for advanced tissue-engineered constructs.

While HH signalling is pivotal in driving lineage commitment, it is insufficient as a solitary driver to fully orchestrate complete intramembranous ossification [84, 85]. Our findings also implicate IGF-II as an effector in later-stage osteogenic differentiation. Aberrant IGF-II signalling in HH-responsive cells severely impairs bone formation in mouse embryos, highlighting the necessity of sustained HH expression to activate IGF-II to complete the differentiation process [86], aligning with our findings. The activation of IGF-II by GLI1 extends to ECM regulation, with a bidirectional interaction between *IGF-II* and *ITGA5* promoting *FN1* expression, forming a positive feedback loop that enhances ECM organisation and induces *RUNX2* expression in hMSCs [87]. Furthermore, IGF-II has been reported to promote osteoblast maturation up to bone mineralisation [88]. We previously demonstrated that topographically-textured microparticles can be used to induce bone regeneration *in vivo*, confirming their osteogenic commitment [15].

To translate these findings, 2PP was used to fabricate GLI1 expression gradients with subcellular resolution [89]. While previous models relied on chemical inducers and genetic modification [90, 91], our platform offers a material-independent, non-genetic alternative with improved scalability and physiological relevance. Li *et al*. engineered ‘sender’ cells to express SHH using a 4-hydroxytamoxifen-inducible system and created open-loop receptor cells by modifying *PTCH1* alleles with a doxycycline (Dox)-inducible promoter [90]. While this 2D system offered valuable insights into HH spatial patterning, its scalability and reproducibility potentially suffered from its high cost [92], and the possibility that Dox may exert off-target effects [93]. Johnson *et al*. developed an SHH morphogen gradient by overlaying a genetically modified epithelial layer producing SHH onto mesenchymal tissues embedded in a hyaluronic acid-collagen gel [91]. However, the model’s reliance on genetic modulation may limit physiological relevance and scalability.

By demonstrating spatial control of GLI1 expression via engineered topographical gradients, evidence is provided herein that mechanical cues can orchestrate cell signalling with precision previously thought possible only through biochemical means, independent of the material used for fabrication. This allows for versatile fabrication using either solvent evaporation oil-in-water emulsion (for scalable, high-throughput microparticle production) or 2PP (for precise, customisable spatial patterning of localised cell behaviour and tissue gradients). It has been demonstrated that GLI1 activity reflects SHH responses in a proportional manner, thereby reflecting subtle changes in SHH concentration without amplification [94]. Our approach enables the modulation of cellular responses by engineered topographical cues in isolation of external biochemical additives. The observed GLI1 gradient spanned approximately 290 μm, aligning with the reported range of SHH morphogen activity of 200 µm [90] and closely parallels the documented SHH signalling range of 300 μm in the limb bud [95]. This spatial control demonstrates that topographical features can function as a precise and reproducible platform for in vitro modelling of spatially patterned stem cell behaviour, with broad relevance to tissue engineering, developmental biology, and drug screening.

### Concluding remarks

We present a versatile, modular microparticle-based platform that enables precise, additive-free control of osteogenic differentiation in hMSCs through engineered 3D topographies. This topography-driven induction, achieved without biochemical additives or genetic manipulation, represents a significant step towards streamlining cell-instructive platform design while avoiding regulatory complexities associated with exogenous growth factors. Transcriptomic profiling underscores the biological potency of topographical programming, and the use of high-resolution 2PP lithography enabled precise engineering of bioinstructive microtopographical cues, supporting reproducible, spatially controlled cell responses. By decoupling topography from soluble signalling, this platform provides a scalable, physiologically relevant system for probing mechanotransduction and differentiation. Beyond osteogenesis, this approach holds promise for broader applications in modelling other developmental or disease contexts where spatial organisation and mechanical inputs play instructive roles, such as vasculogenesis and organoid patterning. The platform is modular and adaptable to different culture formats, which are key characteristics required for scalable and reproducible deployment in translational settings. Collectively, this lays the groundwork for next-generation bioinstructive systems in regenerative medicine, developmental biology, and high-content screening.

## METHODS

### Experimental model and subject details

This study used primary human bone marrow-derived mesenchymal stromal cells obtained from five independent donors representing diverse demographic backgrounds. Three donor lots (designated as donors 1, 2 and 3) were obtained from RoosterBio (RoosterVial™-hBM-1M, MSC-003, RoosterBio Inc., USA), while two additional donor lots (designated as donors 4 and 5) were obtained from Lonza (PT-2501, Lonza, Germany). An overview of donor characteristics is provided in Table S1.

## Methods

### Fabrication of smooth and dimpled microparticles

Poly(D,L-lactic acid) (PLA) microparticles (Ashland Viatel DL 09 E, Mn 56.5 kDa, Mw 111 kDa, IV 0.8–1.0 dL/g) were prepared using a solvent evaporation oil-in-water emulsion technique [15, 17]. For fabricating smooth microparticles, a 20% (*w/v*) solution of PLA in dichloromethane (DCM; ≥99.8%, Thermo Fisher Scientific, USA) was homogenised (Silverson Machines Ltd., UK) at 3800 rpm for 5 min. The homogenised organic phase was then emulsified into 100 mL of an aqueous continuous phase containing 1% (*w/v*) poly(vinyl acetate-co-alcohol) (PVA; MW 13– 23 kDa, #348406, Sigma-Aldrich). The resulting emulsion was stirred at room temperature to facilitate solvent evaporation. Microparticles were collected by centrifugation and washed with deionised water to remove residual PVA. After washing, microparticles were sieved using strainers (40–70 µm) (Greiner bio-one) and then freeze-dried for storage.

For the fabrication of dimpled microparticles, the addition of fusidic acid (FA; 98%, #5552333, Thermo Fisher Scientific, USA) into the organic phase was used to create the topographical patterns. A 30% (*w/v*) ratio of FA to PLA was used, resulting in a total FA/PLA concentration of 10% (*w/v*) in DCM. FA-loaded microparticles were incubated in phosphate-buffered saline (PBS, Gibco) at 37 °C for 7 days for FA release, as previously detailed [15].

### Microparticle size analysis

The particle size distribution of the fabricated microparticles was measured using a laser diffraction particle size analyser (Mastersizer 3000, Malvern Instruments Ltd, UK). Particle size distributions were calculated automatically using the optical Mie model [96] within the Mastersizer 3000 software (v3.71). Each analysis was performed at least three times. The polydispersity index (PDI) of size distribution was determined by dividing the square of the standard deviation by the square of the mean diameter.

### Brunauer-Emmett-Teller (BET) surface area measurements

The surface area of fabricated microparticles was determined as previously described [15]. Krypton (Kr) sorption isotherms were conducted using a Micromeritics ASAP 2420 (Micromeritics, USA) at −196 °C. Approximately 500 mg of each sample was degassed under high vacuum (<10 mTorr) at 37 °C for 48 h to eliminate moisture and adsorbed gases. As Kr solidifies at higher pressures at −196 °C (>1.77 Torr) [97], sorption isotherms were measured over a relative pressure range of 0.10 to 0.65. The specific surface area was calculated from 0.05 to 0.30 relative pressure range using the BET model. Micropore volume was calculated at 2 nm, and limited mesopore volume from 2-20 nm using pore size vs cumulative pore volume using the Derjaguin– Broekhoff–de Boer model [98].

### Atomic force microscopy (AFM) topographical analysis

Atomic force microscopy (AFM) images were acquired on a Bioscope Catalyst AFM (Bruker) mounted on an Eclipse Ti-U (Nikon) inverted optical microscope, with a Nanoscope IV controller, Nanoscope v9.1 software (Bruker), and a ScanAsyst™-Air (Bruker) AFM probe. These silicon nitride probes with an aluminium coating have a nominal spring constant of 0.4 N m^-1^ (Bruker AFM Probes, USA). Imaging was performed using ScanAsyst Peak-Force Tapping mode in air with a scan size of 7 μm at a scan rate of 0.988 Hz. The instrument is periodically calibrated using a grating with 180 nm deep and 10 mm² depressions. Images were processed by applying flattening of first-order using Nanoscope Analysis software (v3.0).

### Primary human mesenchymal stromal cell culture

Primary human bone marrow-derived mesenchymal stromal cells obtained from five independent donors representing diverse demographic backgrounds were used. Three donor lots (designated as donors 1, 2 and 3) were obtained from RoosterBio (RoosterVial™-hBM-1M, MSC-003, RoosterBio Inc., USA), while two additional donor lots (designated as donors 4 and 5) were obtained from Lonza (PT-2501, Lonza, Germany). An overview of the donor characteristics is provided in Table S1. Cells were cultured in Dulbecco’s modified Eagle’s medium (#21969-035, Gibco) supplemented with 1% (*w*/*v*) l-glutamine (Gibco), 1% (*w*/*v*) penicillin-streptomycin (Gibco) and either 10% (*v*/*v*) fetal bovine serum (FBS; Gibco) for routine passaging or 2% FBS (referred to as serum-reduced medium). Each donor batch was maintained as an independent stock and cells were used between passages three and six. Tri-lineage differentiation potential hMSCs was confirmed using StemPro™ differentiation kits (Gibco, UK).

### Microparticles preparation for cell seeding

Microparticles were placed in CELLSTAR^®^ cell-repellent surface 96-well plates (Greiner Bio-One) and sterilised with UV light at 254 nm for 30 min at 4 × 10^4^ mJ. Mass of smooth and dimpled microparticles were calculated to achieve a consistent surface area for cell attachment. Following sterilisation, the microparticles were conditioned in serum-reduced medium (2% FBS) for 1 h. Cells were seeded onto microparticles at a density of 1 × 10^4^ cells/cm² and placed on a plate shaker for 15 min to ensure even distribution. For 2D controls, cells were seeded in tissue culture-treated 96-well plates (CytoOne^®^, Starlab).

### Cell viability and proliferation

Cell viability was assessed three days post-seeding using the Viability/Cytotoxicity Assay Kit for Animal Live & Dead Cells (30002-T, Biotium, UK) according to the manufacturer’s instructions. Briefly, 1 μM calcein-acetoxymethyl (calcein-AM) and 4 μM ethidium homodimer III (EthD-III) were added to each well. Imaging was performed using a ZEISS Cell discoverer 7 imaging system (ZEISS, Germany).

Proliferation was assessed by measuring DNA concentration from cell lysates using the Quant-iT™ PicoGreen^®^ dsDNA Assay Kit (P7589, Invitrogen, USA), following manufacturer’s instructions. Cells were lysed with CelLytic™ M lysis buffer (C2978, Sigma-Aldrich) with two additional freeze-thaw cycles. Fluorescence was measured at λ_exc_/λ_em_ 480/520 nm using a Varioskan™ LUX multimode microplate reader (Thermo Fisher Scientific, USA). DNA content was measured by comparing to a standard curve generated from the supplied standards.

### Scanning electron microscopy (SEM)

Microparticles were directly mounted onto double-sided copper tape and placed on an aluminium pin stub. For imaging cell-microparticle aggregates, cells were fixed after three days of culture using 2.5% (*v/v*) glutaraldehyde (G6257, Sigma-Aldrich). Fixed samples were then dehydrated with a graded ethanol series (10, 25, 50, 80, and 100%; Fisher Chemical). The dehydrated cell aggregates were mounted onto double-sided copper tape (Agar Scientific) and placed on an aluminium pin stub (AGG301, Agar Scientific). Prior to imaging, samples were sputter-coated with an 80% gold/20% palladium alloy using a Q150 S/E/ES Plus sputter coater (Quorum Technologies, UK) under vacuum at 40 mA for 4 min. Scanning electron microscopy (SEM) images were acquired using a Tescan Vega 3 (Tescan, UK) at 5 and 10 kV, as detailed in the figure captions. Dimple sizes were characterised using ImageJ software (v1.53q) by measuring the diameters of a minimum of 250 dimpled microparticles, across ten independent SEM images from five fabricated batches.

### Immunocytochemistry

Cells were fixed with 3.7% (*v/v*) formaldehyde (Thermo Fisher Scientific, USA) in PBS and permeabilised with 0.1% (*v/v*) Triton X-100 (A16046, Fisher Chemicals) in PBS. To block non-specific binding, samples were incubated for 1 h in 1% (*w/v*) bovine serum albumin (BSA; SLCK4263, Sigma-Aldrich) in PBS, supplemented with 10% (*v/v*) normal goat serum (G9023, Sigma-Aldrich) or normal donkey serum (#D9663, Sigma-Aldrich), based on the secondary antibody. Following blocking, cells were incubated with the primary antibody overnight at 4 °C, then the secondary antibody for 2 h. For F-actin staining, cells were stained with ActinGreen™ 488 ReadyProbes™ Reagent (R37110, Invitrogen) and nuclei were counter-stained with NucBlue™ Fixed Cell ReadyProbes™ Reagent (R37606, Invitrogen). Cells were observed using a Zeiss LSM 880 inverted AiryScan confocal microscope (Carl Zeiss, Germany).

The following primary antibodies were used, goat anti-GLI-1 affinity purified polyclonal antibody (1:100; AF3455, R&D systems, RRID: AB_2247710), rabbit anti-osteocalcin (OCN) polyclonal antibody (1:75; AB10911, Millipore, RRID: AB_1587337), goat anti-type I collagen polyclonal antibody (1:200; #1310-01, SouthernBiotech, RRID: AB_2753206) and monoclonal anti-Vinculin (1:70; V4505, Sigma-Aldrich, RRID: AB_477617). All secondary antibodies were obtained from Invitrogen and used at a 1:500 dilution, including donkey anti-goat IgG (H+L) cross-adsorbed, Alexa Fluor™ 594 (A-11058, RRID: AB_2534105), goat anti-rabbit IgG (H+L) cross-adsorbed, Alexa Fluor™ 488 (A-11008, RRID: AB_143165), donkey anti-goat IgG (H+L) highly cross-adsorbed, Alexa Fluor™ Plus 488 (A32814, RRID: AB_2762838), and goat anti-mouse IgG (H + L) cross-adsorbed, Alexa Fluor™ 647 (A-21235, RRID: AB_2535804). Nuclei were counter-stained with NucBlue™ Fixed Cell ReadyProbes™ (R37606, Invitrogen).

### Gene expression analysis using real-time PCR

Total RNA samples were extracted using the RNAqueous™-Micro Kit (AM1931, Thermo Fisher Scientific) with minor modifications to prevent dissolution of microparticles in ethanol. The concentration of RNA in each sample was measured using a NanoDrop™ 1000 Spectrophotometer (Thermo Fisher Scientific). Reverse transcription of RNA was achieved using the iScript™ Select cDNA Synthesis Kit (#1708896, Bio-Rad, USA) following the manufacturer’s protocol. A GS00482 Thermal Cycler (G-STORM, UK) was used following the reaction conditions listed in Table S6. Transcript levels were determined using SsoFast™ EvaGreen^®^ Supermix (#1725201, Bio-Rad), quantified using CFX384 Touch Real-Time PCR Detection System (Bio-Rad) and following the reaction conditions listed in Table S6. Primer sequences are provided in Table 2, and were used at a concentration of 500 nM with an annealing temperature of 64 °C. No template controls (NTC) for each primer set and no reverse-transcriptase controls (NRT) for each sample were included.

**Table 2:**
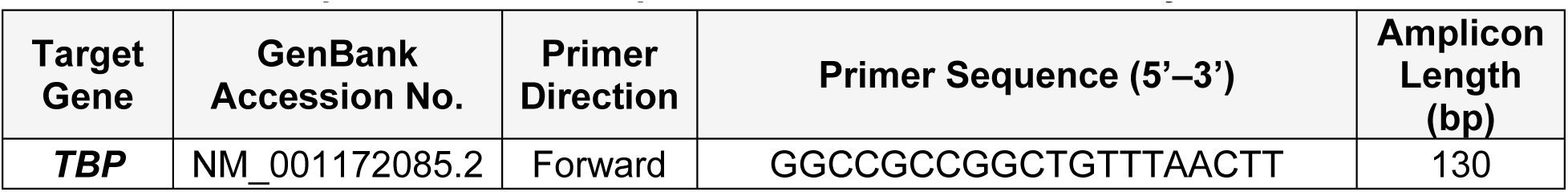
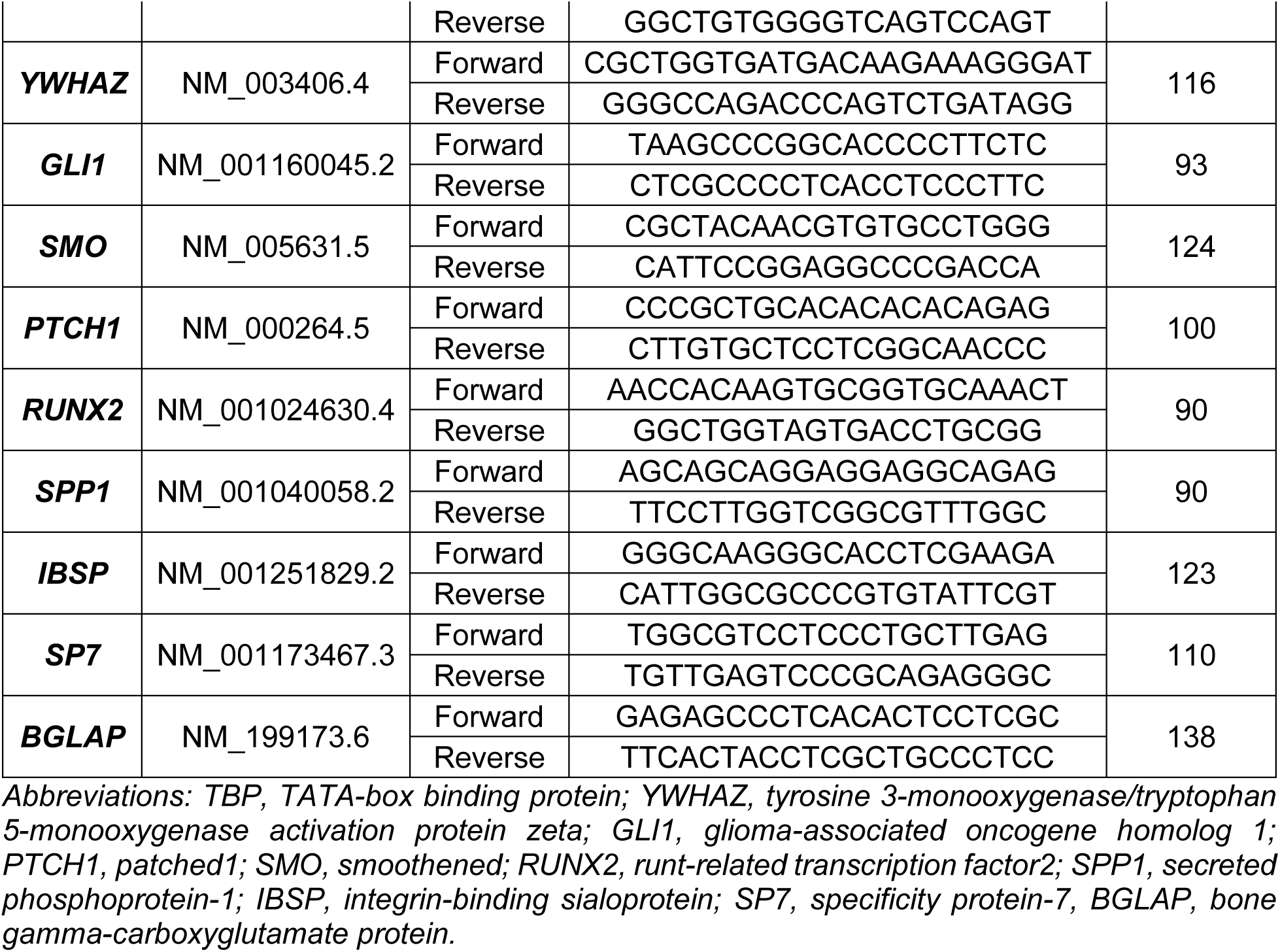
Primer sequences used for quantitative real-time PCR analysis.

Purmorphamine (#130-104-465, StemMACS™ Purmorphamine, Miltenyi Biotec) treatment was used as a positive 2D control for *GLI1*, *SMO*, *PTCH1* and *RUNX2* expression, dissolved in dimethyl sulfoxide (DMSO; 5 mM), and used as 2 μM in serum-reduced medium (2% FBS). Corresponding 2D vehicle-only controls were prepared as 0.06% (*v/v*) DMSO in serum-reduced medium (2% FBS). Cells in 2D culture were treated with osteoinductive media containing dexamethasone (A1007201, StemPro, Gibco, UK) as positive 2D controls for osteoblastic markers, except for *RUNX2*, as dexamethasone has been reported to promote osteogenesis indirectly by inhibiting chondrogenesis rather than directly inducing *RUNX2* expression [2]. Corresponding negative 2D controls were cultured in serum-reduced medium. Relative expression levels of each gene of interest were calculated using the 2^−ΔΔCt^ method [99], normalised to the geometric mean of two housekeeping genes, TATA-box binding protein (*TBP*) and tyrosine 3-monooxygenase/tryptophan 5-monooxygenase activation protein zeta (*YWHAZ*), known for their stable expression in hMSCs during osteogenic differentiation [100, 101]. Results were further normalised to corresponding untreated 2D controls.

### Treatment with KAAD-cyclopamine

HMSCs were cultured in serum-reduced medium for 24 h, then treated with 300 nM KAAD-cyclopamine (ab142146, Abcam), an SMO antagonist. Corresponding 2D and 3D vehicle-only controls, were prepared as 0.06% (*v/v*) DMSO in serum-reduced medium, shown to maintain the surface topographical features intact [17], for 3 and 14 days, with the treatment media refreshed every 2 days. Cells treated with KAAD-cyclopamine will be referred to as KAAD-cyclopamine –treated.

### RNA-Seq and library preparation

hMSCs (N= 3) were cultured on smooth and dimpled microparticles as previously outlined. Total RNA was extracted on days 3 and 14 post-seeding, as detailed above. Total RNA quality was assessed using the Agilent 4200 TapeStation system with RNA ScreenTape assays (G2991BA, Agilent Technologies Inc), and the RNA Integrity Number (RIN) was determined using TapeStation Analysis Software (v5.1; Agilent Technologies). Unmapped paired-end sequences from NovaSeq 6000 sequencer were tested by FastQC (Available from: http://www.bioinformatics.babraham.ac.uk/projects/fastqc/). Sequence adapters were removed, and reads were quality trimmed using Trimmomatic_0.39 [102].

Reads were mapped against the reference human genome (hg38) and counts per gene were calculated using annotation from GENCODE 44 (http://www.gencodegenes.org/) using STAR_2.7.7a [103]. RNA samples from donor 2 (MSCs seeded on untreated dimpled microparticles at day 3) were excluded because the read counts assigned to genes were below our threshold of 1 × 10⁷ [104] (Figure S4). Normalised read counts were averaged for each gene across the conditions of interest; genes with a mean normalised read count < 10 were excluded from further analysis. Normalisation, Principal Components Analysis (PCA), and differentially expressed genes (DEGs) was calculated with DESeq2_1.40.2 [105]. Adjusted *p*-values (*p*_adj_) were corrected for multiple testing (Benjamini and Hochberg method). Heatmaps were drawn with ComplexHeatmap v2.16.0 [106]. Hierarchical clustering was performed on the means of each sample group (log_2_) that had been z-transformed (for each gene the mean set to zero, standard deviation to 1). A threshold of log_2_ fold change > 1 (upregulated) or < −1 (downregulated) and an adjusted *p*-value (*p*_adj_) < 0.05 was applied to identify significantly differentially expressed genes. Gene ontology enrichment was studied using clusterProfiler v4.8.3 [107] and Enrichr v3.2 [108]. Gene enrichment was studied using ReactomePA 1.44.0 [109].

For Ingenuity Pathway Analysis (IPA), a more stringent filter was applied, excluding genes with a mean normalised read count < 50 to enhance confidence in pathway predictions. Canonical pathways and upstream regulators at day 14 post-seeding were identified/predicted using IPA software (QIAGEN). DEGs with a log_2_ fold change > 2 or < −2 and *p*_adj_ < 0.05 were used to perform core analyses. Canonical pathways were ranked based on a –log(*p*-value) threshold > 1.3, which corresponds to *p*_adj_ < 0.05, calculated using Fisher’s Exact Test to determine statistical significance of enrichment [110]. The activation z-score was employed to predict the activation or inhibition of pathways by comparing the observed gene expression patterns against curated information in the IPA Knowledge Base. Pathways were considered significantly impacted based on z-score threshold of > 2, indicating predicted activation, or < –2, indicating predicted inhibition. To explore IGF-II regulatory mechanisms, the Interaction Network tool in IPA was applied to DEGs identified in dimpled versus smooth cultures. Networks were trimmed using IPA’s built-in filtering functions to exclude low-confidence nodes and genes that did not meet differential expression thresholds or were absent from the input dataset. The IGF-II-centred interaction network was graphically simplified using BioRender to enhance visual clarity. The IPA-generated network is shown in Figure S3, with simplified visualisation in Figure 6A.

### Multiplex fluorescent western blotting

After 14 days in culture, cells were lysed using CelLytic™ M lysis buffer (C2978, Sigma-Aldrich) containing 1 mM phenylmethyl sulfonyl fluoride (PMSF; #36978, Thermo Fisher Scientific) and 1 mM ethylenediaminetetraacetic acid (EDTA; #46-034-CI, Corning) at pH 8.0, and 1 μL protease inhibitor cocktail (P8340, Sigma-Aldrich). Following incubation on ice for 20 min with continuous shaking, cell lysates were centrifuged for 15 min at 13,000 *× g*. Total protein concentrations were determined using the Pierce™ Bicinchoninic acid (BCA) Protein Assay Kit (#23227, Thermo Fisher Scientific). Equal amounts of protein were prepared in NuPAGE (4X) lithium dodecyl sulphate sample buffer (NP0007, Invitrogen) with 2-mercaptoethanol as a reducing agent (M3148, Sigma-Aldrich) and denatured at 95 °C for 5 min. Samples were then subjected to SDS-PAGE on a pre-cast NuPAGE™ 4–12% Bis-Tris gel (NP0335B0X, Invitrogen) and transferred to a nitrocellulose membrane using the Trans-Blot Turbo Transfer system (#1704270, Bio-Rad). Membranes were blocked in PBS with 5% non-fat milk and 0.1% Tween-20 for 1 h on a rolling shaker at room temperature and incubated with primary antibodies overnight at 4 °C. Goat anti-IGF-I polyclonal antibody (1:5000; AF-291-SP, R&D Systems, RRID: AB_2122119), mouse anti-IGF-II monoclonal antibody (1:1000; MAB2921, R&D Systems, RRID: AB_2233454) and goat anti-Glyceraldehyde-3-Phosphate Dehydrogenase (GAPDH) polyclonal antibody (1:1000, AF5718, R&D Systems, RRID: AB_2278695) were used simultaneously for primary detection. Secondary detection was performed using IRDye 800CW donkey anti-goat IgG (#926-32214, LI-COR Biosciences, RRID: AB_621846) for IGF-I and GAPDH, detected in the blue channel, and IRDye 680RD donkey anti-mouse IgG (#926-68072, LI-COR Biosciences, RRID: AB_10953628) for IGF-II, detected in the red channel. Fluorescence signals were visualised in a ChemiDoc™ MP Imaging System using Image Lab software (v6.1.0, Bio-Rad).

### Two-photon polymerisation (2PP) lithography

Computer-aided design (CAD) files were created using Autodesk Fusion 360 (v.2.0.19941) and Materialise Magics (v27.0), then exported as Standard Tessellation Language (STL) files. Single hemispherical microstructures (27.5 µm height) with defined dimple diameters (2 or 7 µm) served as building blocks for the assembly of array configurations. Structures were imported into the DeScribe software (v2.7, Nanoscribe GmbH), arranged along the x and y axes to generate the required layouts, and converted to GWL format (Table S7). Direct laser writing was performed using NanoWrite (v1.10.5) on the Photonic Professional GT system (Nanoscribe GmbH, Germany).

To initiate the fabrication process, a droplet of IP-Visio photoresin (Nanoscribe GmbH, Germany) was deposited onto the fused silica substrate coated with indium doped tin oxide (ITO). The substrate was then secured to the sample holder and mounted in a holder compatible with the piezoelectric stage. Arrays were written in a bottom-up sequence, with the first layer adhered directly to the substrate surface using a 25x magnification/0.8 NA microscope objective (0.8 DIC Imm Korr, Carl Zeiss AG).

Uncured resin was removed by alternating washes with propylene glycol monomethyl ether acetate (PGMEA; #484431, Sigma-Aldrich) followed by isopropyl alcohol (IPA; I9030, Sigma-Aldrich). This cycle was repeated until all uncured resin was completely removed, and printed constructs were then allowed to air dry for 3-5 min.

### Post-processing the printed arrays and assessment of GLI1 expression

A two-step process was used to quench resin autofluorescence. First, UV bleaching was performed by exposing printed arrays to UV at 365 nm for 3 h at 4 × 10^4^ mJ (CL-1000, Analytik Jena, US). The sample stage was positioned 5 cm from the UV source and encased in aluminium foil to increase the efficiency of UV exposure using its reflective properties. Afterwards, TrueBlack^®^ Lipofuscin Autofluorescence Quencher (#23007, Biotium) was applied according to the manufacturer’s protocol. Briefly, a 1X TrueBlack solution in 70% ethanol was added to cover the samples for 7 min, followed by a thorough, gentle wash with 1X PBS to remove excess solution. Treated arrays were sterilised and conditioned before cell seeding, as described above.

GLI1 expression was detected by immunostaining after 7 days in culture (See: Immunocytochemistry). Confocal fluorescence microscopy images were processed using ImageJ software (v1.53q). Although TrueBlack^®^ and UV bleaching have been reported to effectively quench major autofluorescent structures while preserving immunofluorescence signals [111, 112] autofluorescence in confocal imaging remained a significant challenge in dual-topography arrays. Therefore, for the 3D arrays, a sliding paraboloid background subtraction (rolling ball radius = 0.6 px) was applied to the red channel in the sum-projected confocal images. This method was selected to accommodate local intensity variations and complex geometries [113], ensuring accurate background estimation without over-subtraction near high-intensity features. The rolling ball radius was determined based on the scale of background noise relative to image features and was applied consistently across all red fluorescence channel images. Using different methods for 3D and 2D samples was necessary to ensure optimal background correction for each sample type. For corresponding 2D controls, ImageJ’s Math>Subtract method (8 px) was applied to the red channel in the sum-projected confocal images.

### Statistical analysis

Statistical analysis was performed using GraphPad Prism 9.3.1 (GraphPad Software Inc., USA). Data distribution was first assessed using Shapiro-Wilk and Kolmogorov-Smirnov normality tests. Parametric one-way or two-way ANOVA with Tukey’s or Dunnett’s *post-hoc* tests were used where appropriate. For experiments with N= 3 or datasets that did not meet normality assumptions, non-parametric Freidman test with Dunn’s multiple comparisons test for more than two groups. Data are presented as mean ± standard deviation (SD), with *p* < 0.05 considered the threshold for statistical significance.

## Acknowledgements

FG is supported by a scholarship from Kuwait University. MA acknowledges support by the Academy of Medical Sciences Springboard Scheme [SBF008\1057]. We acknowledge Dr Rachel Saunders (University of Manchester) for assistance with 2PP printing, Dr. Steven Marsden for assistance with AFM microscopy, Dr David Spiller for assistance with confocal microscopy, Dr. Jamie Tibble and Mrs Shahla Khan for assistance with Mastersizer. The Bioimaging Core Facility AFMs used in this study were purchased with grants from BBSRC, Wellcome, Walgreen Boots Alliance and the University of Manchester Strategic Fund. This work was also supported by the Henry Royce Institute for Advanced Materials, funded through EPSRC grants EP/R00661X/1, EP/S019367/1, EP/P025021/1 and EP/P025498/.

## Supplementary information

Supplementary material (attached)

