## Supplementary data for "A Modular Bioinstructive Platform for Additive-Free, Topography-Driven Stem Cell Differentiation and Patterning"

**Figure S1**

**Figure S2**

**Figure S3**

**Figure S4**

**Table S1**

**Table S2**

**Table S3**

**Table S4**

**Table S5**

**Table S6**

**Table S7**

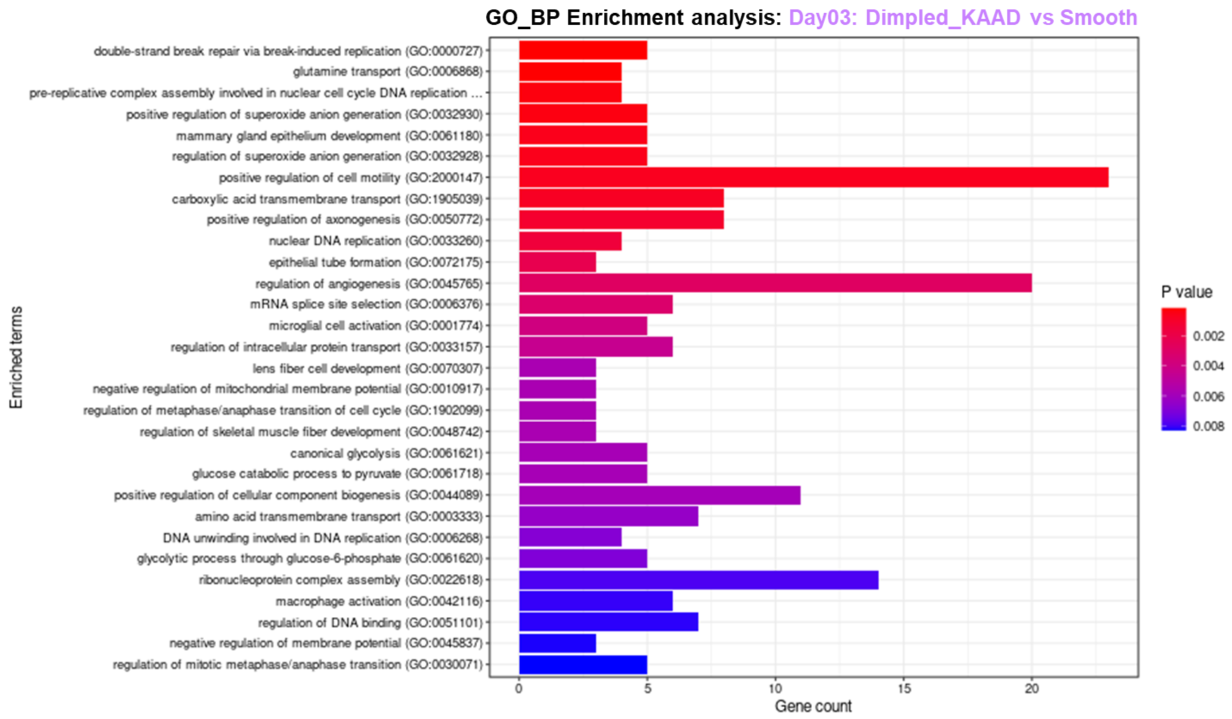


**Figure S1: Gene ontology enrichment analysis for biological processes in hMSCs cultured on dimpled treated with KAAD-cyclopamine versus smooth after 3 days in culture, performed with Enrichr.** Gene count is indicated on the x-axis. Colour indicates Benjamini-Hochberg adjusted *p* value.

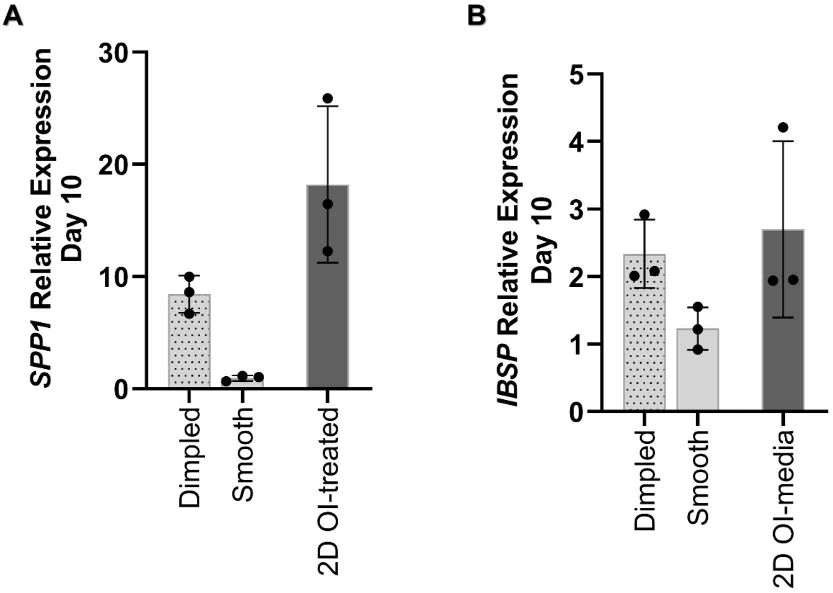


**Figure S2: Relative quantitative real-time PCR analysis of gene expression at day 10, in serum-reduced media, relative to 2D controls: *SPP1* (A) and *IBSP* (B).** A 2D positive control treated with osteoinductive medium is included as reference. Statistical analysis was conducted using Freidman test (N= 3 donors).

*Abbreviations: SPP1, secreted phosphoprotein 1; IBSP, integrin-binding sialoprotein; OI media, Osteoinductive media.*

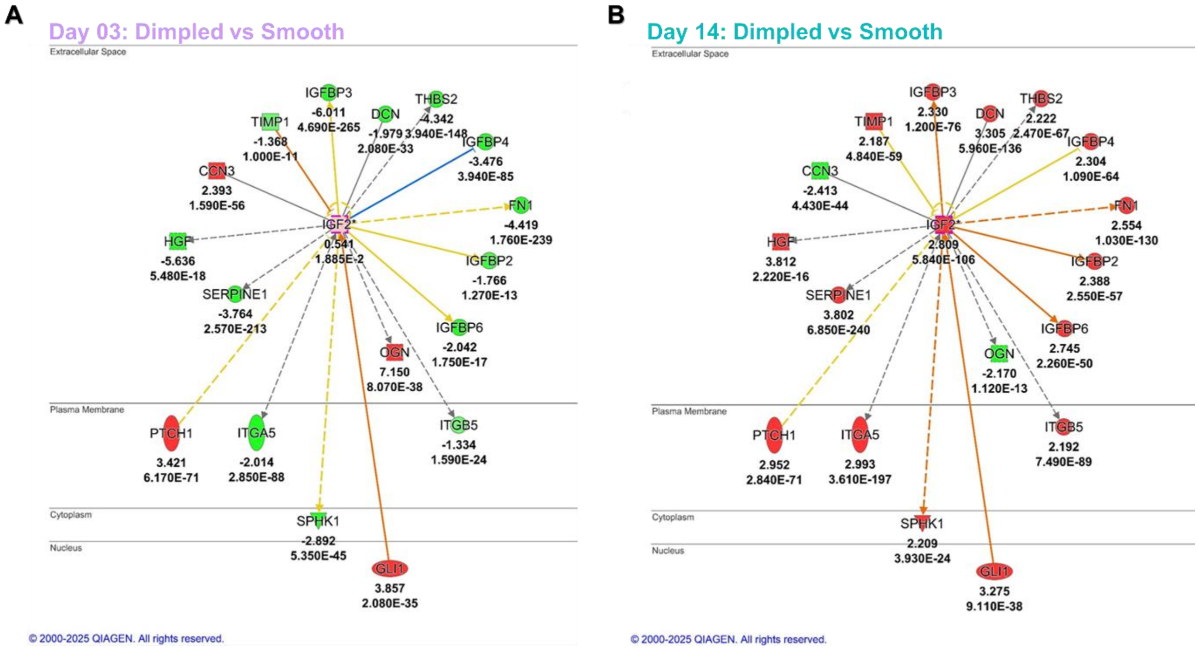


**Figure S3: Network analysis of the upstream regulatory network of IGF-II generated using IPA based on DEGs in dimpled versus smooth samples, highlighting transcriptional regulators, growth factors, and transmembrane receptors associated with IGF-II signalling, including connections to GLI1.** The network was generated by overlaying the DEGs at day 3 (A) and 14 (B) post-seeding. Nodes are colour-coded to represent expression levels: upregulated (red) and downregulated (green). Edges represent predicted relationships: activation (orange), findings inconsistent with expected relationships (yellow), and undefined effects (gray). Nodes not passing the statistical cut-off or not present in the DEG dataset were excluded from the visualisation using the Trim function in Ingenuity Pathway Analysis (IPA) to improve clarity.

*Abbreviations: FN1, Fibronectin 1; PTCH1; Patched 1; GLI1, Glioma Associated Oncogene Homolog; ITGA5, Integrin Subunit Alpha 5; IGF2, Insulin-like Growth Factor 2; IGFBP, Insulin-like Growth Factor Binding Protein; THBS2, Thrombospondin 2; TIMP1, TIMP Metallopeptidase Inhibitor 1; SPHK1, Sphingosine Kinase 1; DCN, Decorin; HGF, Hepatocyte Growth Factor; SERPINE1, Serpin Family E Member 1; CCN3, Cellular Communication Network Factor 3; OGN, Osteoglycin; GAPDH, Glyceraldehyde-3-Phosphate Dehydrogenase; AU, Arbitrary Unit.*

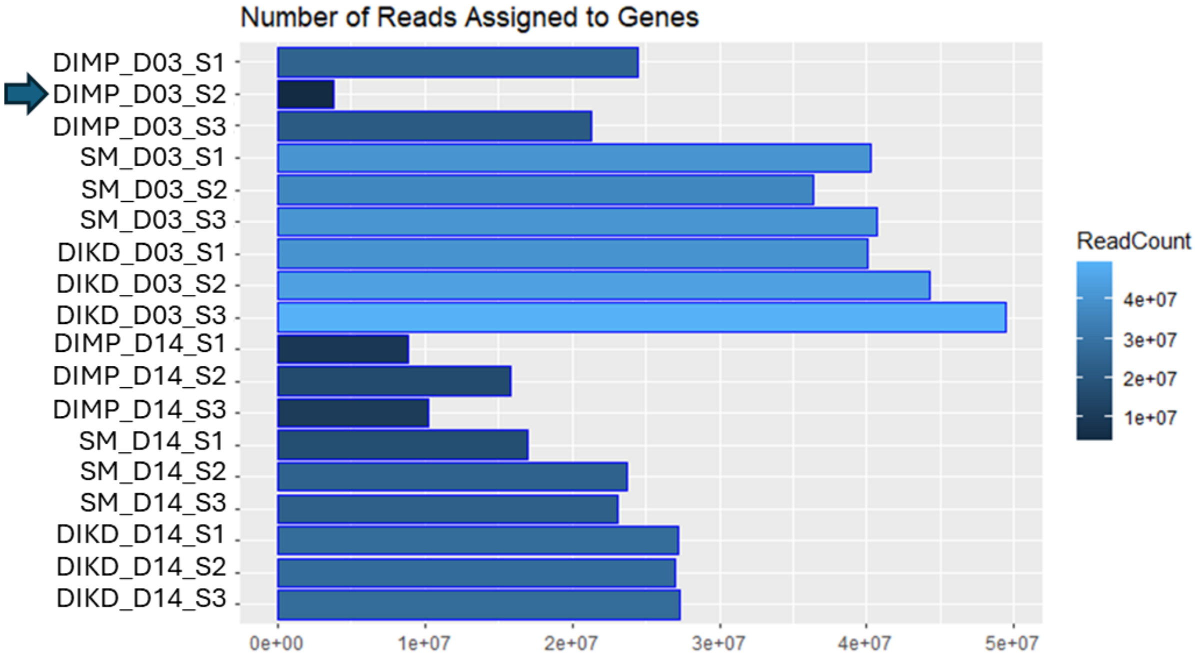


**Figure S4: Quality control analysis of RNA sequencing data for hMSCs cultured on dimpled and smooth microparticles at day 3 and 14 post-seeding.** Bar plot showing the number of reads assigned to genes for each sample, with darker shades of blue representing higher read counts. Arrow indicates DIMO_D03_S2 which corresponds to human MSCs Lot#310307 (Donor 2) seeded on dimpled microparticles at day 3 post-seeding, which was excluded from further analysis at day 3 due to low-quality metrics.

*Abbreviations: DIMP; Dimpled, SM, Smooth; DIKD, Dimpled treated with KAAD-cyclopamine; D03, Day 3 post-seeding; D14, Day 14 post-seeding; S1, donor lot#310305; S2, donor lot#310307; S3, donor lot#310310.*

**Table S1: Overview of the demographic characteristics of hMSCs donors used in this study.**

| Donor ID | Supplier | Cat number | Lot number | Age | Sex | Race |
|---|---|---|---|---|---|---|
| 1 | RoosterBio | MSC-003 | 310305 | 20 | Female | African American |
| 2 | RoosterBio | MSC-003 | 310307 | 19 | Male | Eritrean/East African |
| 3 | RoosterBio | MSC-003 | 310310 | 25 | Male | Caucasian |
| 4 | Lonza | PT_2501 | 0000-491129 | 34 | Female | Caucasian |
| 5 | Lonza | PT_2501 | 0000-411107 | 21 | Female | African American |

**Table S2: Top 30 differentially expressed genes identified in hMSCs cultured on dimpled versus smooth microparticles at day 3 post-seeding.**

| Top 30 differentially expressed genes upregulated_Day3: Dimpled vs Smooth | | | | | | | |
|---|---|---|---|---|---|---|---|
| **Gene symbol** | **$log_2$ FC** | **$p_{adj}$-value** | **Functional category** | **Gene symbol** | **$log_2$ FC** | **$p_{adj}$-value** | **Functional category** |
| *KRT75* | 13.50 | $4.13 \times 10^{-28}$ | Cytoskeleton organisation | *UBBP3* | 11.70 | $1.82 \times 10^{-21}$ | Pseudogene |
| *ENSG00000225991* | 12.93 | $2.26 \times 10^{-26}$ | Pseudogene | *LINC00958* | 11.62 | $2.08 \times 10^{-19}$ | Long non-coding RNA |
| *ENSG00000179131* | 12.47 | $3.24 \times 10^{-24}$ | Pseudogene | *ENSG00000216412* | 11.60 | $2.77 \times 10^{-21}$ | Pseudogene |
| *ENSG00000234017* | 12.18 | $1.70 \times 10^{-23}$ | Pseudogene | *GATA4* | 11.47 | $3.97 \times 10^{-21}$ | Musculoskeletal development and bone maintenance |
| *SLC25A5P5* | 12.07 | $3.41 \times 10^{-23}$ | Pseudogene | *RPL27AP3* | 11.47 | $1.74 \times 10^{-20}$ | Pseudogene |
| *ASTN1* | 11.98 | $3.87 \times 10^{-21}$ | Cytoskeleton organisation | *ENSG00000248448* | 11.46 | $1.15 \times 10^{-20}$ | Pseudogene |
| *FAM95A* | 11.94 | $5.62 \times 10^{-21}$ | Maintenance of genomic stability | *FXYD6* | 11.26 | $4.69 \times 10^{-19}$ | Ion transport and homeostasis |
| *ENSG00000224282* | 11.91 | $3.77 \times 10^{-22}$ | Pseudogene | *EDN2* | 11.16 | $2.03 \times 10^{-18}$ | Stress response and proliferation |
| *BLACAT1* | 11.89 | $2.82 \times 10^{-14}$ | Transcriptional regulation and stem cell maintenance | *ENSG00000213432* | 11.15 | $2.17 \times 10^{-19}$ | Pseudogene |
| *ENSG00000235817* | 11.85 | $3.72 \times 10^{-22}$ | Pseudogene | *ENSG00000249199* | 11.09 | $1.02 \times 10^{-16}$ | Long non-coding RNA |
| *SOX2* | 11.83 | $2.11 \times 10^{-20}$ | Musculoskeletal development and bone maintenance | *CDH18* | 11.08 | $1.00 \times 10^{-18}$ | Metabolism and biosynthesis |
| *KLHDC8A* | 11.80 | $3.63 \times 10^{-22}$ | Stress response and proliferation | *CALN1* | 11.05 | $3.86 \times 10^{-18}$ | Metabolism and biosynthesis |
| *TUBA3C* | 11.80 | $1.38 \times 10^{-43}$ | Cytoskeleton organisation | *DPYSL5* | 11.02 | $1.60 \times 10^{-19}$ | Cytoskeleton organisation |
| *ENSG00000213495* | 11.77 | $1.05 \times 10^{-21}$ | Pseudogene | *SCARA5* | 11.02 | $2.45 \times 10^{-19}$ | Musculoskeletal development and bone maintenance |
| *RPS11* | 11.71 | $1.33 \times 10^{-21}$ | Metabolism and biosynthesis | *ENSG00000229753* | 10.99 | $3.08 \times 10^{-19}$ | Pseudogene |
| **Top 30 differentially expressed genes downregulated_Day3: Dimpled vs Smooth** | | | | | | | |

| Gene symbol | $\log_2$ FC | $p_{adj}$-value | Functional category | Gene symbol | $\log_2$ FC | $p_{adj}$-value | Functional category |
|---|---|---|---|---|---|---|---|
| *KCTD4* | -11.07 | 3.10 x$10^{-13}$ | Ion transport and homeostasis | *C11orf87* | -7.50 | 1.28 x$10^{-45}$ | Primarily expressed in brain tissue |
| *AFAP1-AS1* | -10.44 | 1.76 x$10^{-09}$ | Cytoskeleton organisation | *ENSG00000261379* | -7.46 | 1.51 x$10^{-04}$ | Long non-coding RNA |
| *IFNE* | -9.80 | 3.94 x$10^{-10}$ | Immune and inflammatory responses | *MEG9* | -7.34 | 7.53 x$10^{-69}$ | Transcriptional regulation and stem cell maintenance |
| *ENSG00000286518* | -9.54 | 5.05 x$10^{-09}$ | Pseudogene | *ADCY4* | -7.33 | 1.25 x$10^{-34}$ | Metabolism and biosynthesis |
| *ENSG00000272574* | -9.38 | 8.41 x$10^{-04}$ | Pseudogene | *CARMN* | -7.31 | 2.85 x$10^{-131}$ | Transcriptional regulation and stem cell maintenance |
| *SLC7A14* | -9.37 | 2.39 x$10^{-09}$ | Ion transport and homeostasis | *EPGN* | -7.22 | 9.19 x$10^{-30}$ | Stress response and proliferation |
| *ENSG00000257221* | -8.56 | 2.49 x$10^{-08}$ | Long non-coding RNA | *RFX8* | -6.96 | 7.16 x$10^{-69}$ | Transcriptional regulation and stem cell maintenance |
| *MEG3* | -8.46 | 3.91 x$10^{-166}$ | Transcriptional regulation and stem cell maintenance | *IL6* | -6.84 | 2.53 x$10^{-54}$ | Immune and inflammatory responses |
| *LMO7DN* | -8.22 | 1.23 x$10^{-07}$ | Transcriptional regulation and stem cell maintenance | *ENSG00000224431* | -6.83 | 1.19 x$10^{-08}$ | Pseudogene |
| *GARIN1A* | -8.08 | 2.29 x$10^{-24}$ | Cytoskeleton organisation | *ELN-AS1* | -6.81 | 1.44 x$10^{-12}$ | Long non-coding RNA |
| *ENSG00000279822* | -7.94 | 4.91 x$10^{-04}$ | Long intergenic non-coding RNA | *ABI3BP* | -6.70 | 0 | Cytoskeleton organisation |
| *MEG8* | -7.90 | 5.25 x$10^{-18}$ | Transcriptional regulation and stem cell maintenance | *GDNF* | -6.61 | 2.73 x$10^{-124}$ | Stress response and proliferation |
| *FLG* | -7.60 | 6.80 x$10^{-111}$ | Structural proteins and barrier formation | *ENSG00000261786* | -6.49 | 1.60 x$10^{-124}$ | Long intergenic non-coding RNA |
| *PLCH2* | -7.55 | 4.04 x$10^{-21}$ | Ion transport and homeostasis | *ENSG00000280435* | -6.49 | 1.02 x$10^{-32}$ | Long intergenic non-coding RNA |
| *MIR137HG* | -7.55 | 7.65 x$10^{-32}$ | Transcriptional regulation and stem cell maintenance | *ERMN* | -6.48 | 1.60 x$10^{-03}$ | Cytoskeleton organisation |

*Abbreviations: ABI3BP, ABI family member 3 binding protein; ADCY4, Adenylate Cyclase 4; AFAP1-AS1, Actin Filament Associated Protein 1 Antisense RNA 1; ASTN1, Astrotactin 1; CALN1, Calneuron 1; CARMN, Cardiac Mesoderm Enhancer-Associated Non-Coding RNA; CDH18, Cadherin 18; DPYSL5, Dihydropyrimidinase-Like 5; EDN2, Endothelin 2; ELN-AS1, ELN Antisense RNA 1; EPGN, Epithelial Mitogen; ERMN, Ermin; FLG, Filaggrin; FXYD6, FXYD Domain Containing Ion Transport Regulator 6; GATA4, GATA Binding Protein 4; GDNF, Glial Cell Line-Derived Neurotrophic Factor; IFNE, Interferon Epsilon; IL6, Interleukin 6; KLHDC8A, Kelch Domain Containing 8A; KRT75, Keratin 75; LINC00958, Long Intergenic Non-Protein Coding RNA 958; LMO7DN, LIM Domain 7 Downstream Neighbor; MEG3, Maternally Expressed Gene 3; MEG8, Maternally Expressed Gene 8; MEG9, Maternally Expressed Gene 9; PLCH2, Phospholipase C Eta 2; RFX8, Regulatory Factor X8; SCARA5, Scavenger Receptor Class A Member 5; SLC7A14, Solute Carrier Family 7 Member 14; SOX2, SRY-Box Transcription Factor 2; TUBA3C, Tubulin Alpha 3C; UBBP3, Ubiquitin B Pseudogene 3.*

**Table S3: Top 30 differentially expressed genes identified in hMSCs cultured on dimpled versus smooth microparticles at day 14 post-seeding.**

| Top 30 differentially expressed genes upregulated_Day14: Dimpled vs Smooth | | | | | | | |
|---|---|---|---|---|---|---|---|
| **Gene symbol** | **$log_2$ FC** | **$p_{adj}$-value** | **Functional category** | **Gene symbol** | **$log_2$ FC** | **$p_{adj}$-value** | **Functional category** |
| *SOSTDC1* | 7.89 | $1.60 \times 10^{-16}$ | Musculoskeletal development and bone maintenance | *CNTNAP3B* | 4.65 | $3.03 \times 10^{-16}$ | Cell adhesion and neuronal signalling |
| *C7* | 6.82 | $2.83 \times 10^{-73}$ | Immune and inflammatory responses | *SLC38A5* | 4.63 | $8.22 \times 10^{-30}$ | Metabolism and biosynthesis |
| *CHI3L1* | 6.50 | $2.45 \times 10^{-92}$ | Immune and inflammatory responses | *PCSK1* | 4.49 | $1.48 \times 10^{-17}$ | Metabolism and biosynthesis |
| *APCDD1L* | 5.87 | $1.36 \times 10^{-171}$ | Musculoskeletal development and bone maintenance | *ACKR1* | 4.48 | $3.37 \times 10^{-08}$ | Immune and inflammatory responses |
| *INSC* | 5.86 | $1.05 \times 10^{-17}$ | Transcriptional regulation and stem cell maintenance | *KRTAP1-5* | 4.47 | $6.28 \times 10^{-10}$ | Structural proteins and barrier formation |
| *FGF5* | 5.48 | $1.61 \times 10^{-37}$ | Musculoskeletal development and bone maintenance | *SERPINE2* | 4.47 | 0 | Musculoskeletal development and bone maintenance |
| *CXCL5* | 5.43 | $4.87 \times 10^{-12}$ | Immune and inflammatory responses | *POU3F3* | 4.43 | $7.72 \times 10^{-32}$ | Musculoskeletal development and bone maintenance |
| *NTM* | 5.27 | $1.04 \times 10^{-28}$ | Transcriptional regulation and stem cell maintenance | *ABLIM3* | 4.43 | $1.32 \times 10^{-130}$ | Cytoskeleton organisation |
| *AKR1C2* | 5.23 | $3.98 \times 10^{-69}$ | Musculoskeletal development and bone maintenance | *CADM3* | 4.41 | $5.76 \times 10^{-208}$ | Cytoskeleton organisation |
| *CXCL1* | 4.98 | $2.69 \times 10^{-27}$ | Immune and inflammatory responses | *SEC14L2* | 4.40 | $3.19 \times 10^{-74}$ | Metabolism and biosynthesis |
| *GALNT5* | 4.90 | $2.89 \times 10^{-139}$ | Metabolism and biosynthesis | *ADGRL4* | 4.39 | $6.24 \times 10^{-25}$ | Musculoskeletal development and bone maintenance |
| *HAS1* | 4.89 | $3.11 \times 10^{-14}$ | Musculoskeletal development and bone maintenance | *AKR1C1* | 4.35 | $2.71 \times 10^{-35}$ | Musculoskeletal development and bone maintenance |

| | | | | | | | |
|---|---|---|---|---|---|---|---|
| *CXCL8* | 4.78 | 4.92 x10$^{-09}$ | Immune and inflammatory responses | *CPAMD8* | 4.32 | 1.37 x10$^{-37}$ | Immune and inflammatory responses |
| *CDKN2B* | 4.75 | 3.15 x10$^{-29}$ | Stress response and proliferation | *COL10A1* | 4.32 | 4.56 x10$^{-23}$ | Musculoskeletal development and bone maintenance |
| *SLC14A1* | 4.66 | 3.59 x10$^{-62}$ | Ion transport and homeostasis | *STEAP1B* | 4.30 | 2.02 x10$^{-28}$ | Ion transport and homeostasis |
| **Top 30 differentially expressed genes downregulated_Day14: Dimpled vs Smooth** | | | | | | | |
| **Gene symbol** | **log$_2$ FC** | **$p_{adj}$-value** | **Functional category** | **Gene symbol** | **log$_2$ FC** | **$p_{adj}$-value** | **Functional category** |
| *CNMD* | -9.29 | 4.54 x10$^{-13}$ | Musculoskeletal development and bone maintenance | *MAMDC4* | -2.42 | 2.81 x10$^{-07}$ | Structural proteins and barrier formation |
| *SFRP2* | -5.07 | 1.15 x10$^{-29}$ | Musculoskeletal development and bone maintenance | *CCN3* | -2.41 | 9.13 x10$^{-42}$ | Musculoskeletal development and bone maintenance |
| *PGM5* | -4.18 | 3.00 x10$^{-97}$ | Musculoskeletal development and bone maintenance | *MARCHF10* | -2.34 | 7.60 x10$^{-10}$ | Stress response and proliferation |
| *RSPO2* | -3.87 | 3.86 x10$^{-48}$ | Musculoskeletal development and bone maintenance | *SPX* | -2.34 | 3.22 x10$^{-16}$ | Metabolism and biosynthesis |
| *RTEL1-TNFRSF6B* | -3.61 | 7.89 x10$^{-10}$ | Maintenance of genomic stability | *TNMD* | -2.33 | 7.61 x10$^{-40}$ | Musculoskeletal development and bone maintenance |
| *TGM1* | -3.52 | 8.35 x10$^{-11}$ | Structural proteins and barrier formation | *ENSG00000262877* | -2.30 | 7.67 x10$^{-05}$ | Long intergenic non-coding RNA |
| *FGF18* | -2.78 | 1.92 x10$^{-13}$ | Musculoskeletal development and bone maintenance | *OLR1* | -2.26 | 8.16 x10$^{-28}$ | Immune and inflammatory responses |
| *COLCA1* | -2.68 | 5.08 x10$^{-08}$ | Transcriptional regulation and stem cell maintenance | *DRP2* | -2.23 | 1.70 x10$^{-09}$ | Cytoskeleton organisation |
| *CCDC78* | -2.67 | 9.33 x10$^{-03}$ | Musculoskeletal development and bone maintenance | *ANKRD2* | -2.22 | 1.12 x10$^{-04}$ | Musculoskeletal development and bone maintenance |
| *CDK5RAP3* | -2.61 | 5.29 x10$^{-09}$ | Stress response and proliferation | *AIF1L* | -2.20 | 8.64 x10$^{-70}$ | Cytoskeleton organisation |

| | | | | | | | |
|---|---|---|---|---|---|---|---|
| *CDHR1* | -2.60 | 7.98 $\times10^{-19}$ | Structural proteins and barrier formation | *SPARCL1* | -2.20 | 1.88 $\times10^{-09}$ | Musculoskeletal development and bone maintenance |
| *MIR503HG* | -2.58 | 1.70 $\times10^{-10}$ | Transcriptional regulation and stem cell maintenance | *PPP1R1B* | -2.18 | 3.66 $\times10^{-13}$ | Stress response and proliferation |
| *NPNT* | -2.56 | 5.64 $\times10^{-20}$ | Musculoskeletal development and bone maintenance | *TRPC6* | -2.18 | 6.38 $\times10^{-16}$ | Ion transport and homeostasis |
| *NPPB* | -2.49 | 1.09 $\times10^{-20}$ | Stress response and proliferation | *ADAM33* | -2.18 | 3.73 $\times10^{-33}$ | Musculoskeletal development and bone maintenance |
| *LTB4R* | -2.48 | 8.15 $\times10^{-19}$ | Immune and inflammatory responses | *OGN* | -2.17 | 4.13 $\times10^{-12}$ | Musculoskeletal development and bone maintenance |

*Abbreviations: ABLIM3, Actin Binding LIM Protein Family Member 3; ACKR1, Atypical Chemokine Receptor 1; ADAM33, ADAM Metallopeptidase Domain 33; ADGRL4, Adhesion G Protein-Coupled Receptor L4; AIF1L, Allograft Inflammatory Factor 1 Like; AKR1C1, Aldo-Keto Reductase Family 1 Member C1; AKR1C2, Aldo-Keto Reductase Family 1 Member C2; APCDD1L, APC Down-Regulated 1 Like; CADM3, Cell Adhesion Molecule 3; CCDC78, Coiled-Coil Domain Containing 78; CDHR1, Cadherin Related Family Member 1; CDK5RAP3, CDK5 Regulatory Subunit Associated Protein 3; CDKN2B, Cyclin Dependent Kinase Inhibitor 2B; CHI3L1, Chitinase 3 Like 1; CNMD, Chondromodulin; COL10A1, Collagen Type X Alpha 1 Chain; COLCA1, Colorectal Cancer Associated 1; CPAMD8, Complement Component 4 Binding Protein-Like; CXCL1, C-X-C Motif Chemokine Ligand 1; CXCL5, C-X-C Motif Chemokine Ligand 5; CXCL8, C-X-C Motif Chemokine Ligand 8; FGF18, Fibroblast Growth Factor 18; FGF5, Fibroblast Growth Factor 5; GALNT5, Polypeptide N-Acetylgalactosaminyltransferase 5; HAS1, Hyaluronan Synthase 1; IL6, Interleukin 6; INSC, Inscuteable Spindle Orientation Adaptor Protein; KRTAP1-5, Keratin Associated Protein 1-5; LTB4R, Leukotriene B4 Receptor; MAMDC4, MAM Domain Containing 4; MARCHF10, Membrane Associated Ring-CH-Type Finger 10; MIR503HG, MIR503 Host Gene; NPPB, B-Type Natriuretic Peptide; NPNT, Nephronectin; NTM, Neurotrimin; OGN, Osteoglycin; OLR1, Oxidised Low-Density Lipoprotein Receptor 1; PCSK1, Proprotein Convertase Subtilisin/Kexin Type 1; PGM5, Phosphoglucomutase 5; POU3F3, POU Class 3 Homeobox 3; PPP1R1B, Protein Phosphatase 1 Regulatory Inhibitor Subunit 1B; RSPO2, R-spondin 2; SEC14L2, SEC14 Like Lipid Binding 2; SERPINE2, Serpin Family E Member 2; SFRP2, Secreted Frizzled-Related Protein 2; SLC14A1, Solute Carrier Family 14 Member 1; SLC38A5, Solute Carrier Family 38 Member 5; SPARCL1, SPARC Like 1; SPX, Spexin Hormone; STEAP1B, STEAP Family Member 1B; TGM1, Transglutaminase 1; TNMD, Tenomodulin; TRPC6, Transient Receptor Potential Cation Channel Subfamily C Member 6.*

**Table S4: Summary of enriched Gene Ontology (GO) biological processes (BP) in hMSCs cultured on dimpled versus smooth microparticles at days 3 and 14 post-seeding.**

| Day 03_Dimpled vs Smooth | | | |
|---|---|---|---|
| **Biological process** | **Overlap ratio** | **$p_{adj}$-value** | **Molecules** |
| Extracellular matrix organization (GO:0030198) | 78/300 | $4.54x10^{-31}$ | *COLGALT2;SPARC;COL16A1;COL14A1;ITGB4;LAMC3;ELN;COL12A1;SERPINE1;LOXL3;TNC;CTSV;LAMC1;FGF2;LOXL2;ADAMTSL1;COMP;ADAMTS2;ADAMTS1;CREB3L1;ADAMTSL3;CTSK;TIMP2;CYP1B1;QSOX1;ADAMTS7;POSTN;MMP2;P3H2;BGN;HSPG2;GREM1;MMP14;VCAN;COL2A1;COL4A2;LOX;COL4A1;COL6A2;PXDN;ADAM12;COL6A1;COL8A2;COL8A1;COL4A5;COL6A3;ITGA5;CD44;LAMA5;COL13A1;HTRA1;PDGFA;ADAMTS12;NID2;THBS1;SCUBE3;ACAN;ADAMTS14;FLRT2;HAS1;HAS2;LAMB3;LAMB2;LUM;COL22A1;FN1;COL1A1;COL1A2;COL5A1;COL5A3;ITGA10;COL5A2;ITGA11;COL9A3;COL9A2;TGFBI;RECK;FBN1* |
| Extracellular structure organization (GO:0043062) | 61/216 | $3.05x10^{-26}$ | *SPARC;COL16A1;COL14A1;ITGB4;LAMC3;ELN;SERPINE1;TNC;LAMC1;FGF2;ADAMTSL1;COMP;ADAMTS2;ADAMTS1;ADAMTSL3;ADAMTS7;POSTN;MMP2;BGN;HSPG2;MMP14;VCAN;COL2A1;COL4A2;LOX;COL4A1;COL6A2;PXDN;ADAM12;COL6A1;COL8A2;COL8A1;COL4A5;COL6A3;ITGA5;CD44;LAMA5;COL13A1;PDGFA;ADAMTS12;NID2;THBS1;ACAN;ADAMTS14;LAMB3;LAMB2;LUM;COL22A1;FN1;COL1A1;COL1A2;COL5A1;COL5A3;ITGA10;COL5A2;ITGA11;COL9A3;COL9A2;TGFBI;RECK;FBN1* |
| External encapsulating structure organization (GO:0045229) | 61/217 | $3.05x10^{-26}$ | SPARC;COL16A1;COL14A1;ITGB4;LAMC3;ELN;SERPINE1;TNC;LAMC1;FGF2;ADAMTSL1;COMP;ADAMTS2;ADAMTS1;ADAMTSL3;ADAMTS7;POSTN;MMP2;BGN;HSPG2;MMP14;VCAN;COL2A1;COL4A2;LOX;COL4A1;COL6A2;PXDN;ADAM12;COL6A1;COL8A2;COL8A1;COL4A5;COL6A3;ITGA5;CD44;LAMA5;COL13A1;PDGFA;ADAMTS12;NID2;THBS1;ACAN;ADAMTS14;LAMB3;LAMB2;LUM;COL22A1;FN1;COL1A1;COL1A2;COL5A1;COL5A3;ITGA10;COL5A2;ITGA11;COL9A3;COL9A2;TGFBI;RECK;FBN1 |
| Collagen fibril organization (GO:0030199) | 34/89 | $9.62x10^{-19}$ | *COLGALT2;COL16A1;COL13A1;COL14A1;ITGB4;COL12A1;LOXL3;LOXL2;ADAMTS2;ADAMTS14;CYP1B1;LAMB3;LUM;COL22A1;P3H2;GREM1;COL1A1;COL2A1;COL1A2;COL4A2;LOX;COL5A1;COL4A1;COL6A2;COL5A* |

| | | | |
|---|---|---|---|
| | | | *3;COL6A1;PXDN;COL5A2;COL8A2;COL4A5;COL6A3;COL8A1;COL9A3;COL9A2* |
| Skeletal system development (GO:0001501) | 29/158 | $5.59x10^{-07}$ | *PKDCC;COL13A1;NPR3;CHRD;XYLT1;TNFRSF11B;PRELP;PKD1;ETS2;PAPSS2;COMP;ACAN;CREB3L2;ALX4;SOX9;TGM2;ZBTB16;WNT5A;SULF1;COL1A1;MMP14;VCAN;COL1A2;COL2A1;EVC;COL9A2;FGFR2;CD44;FBN1* |
| Sprouting angiogenesis (GO:0002040) | 15/52 | $7.86x10^{-06}$ | *SEMA5A;CCBE1;VEGFC;PARVA;VEGFD;FGF2;THBS1;LOXL2;VEGFA;GREM1;EFNB2;BMP4;BMPER;RECK;ENG* |
| **Day 14_Dimpled vs Smooth** | | | |
| **Biological process** | **Overlap ratio** | **$p_{adj}$-value** | **Molecules** |
| Extracellular matrix organization (GO:0030198) | 74/300 | $1.09x10^{-27}$ | *VIT;ITGB1;COL18A1;ITGB5;LAMC3;ELN;SERPINE1;LOXL3;TNC;LAMC2;ICAM5;LAMC1;ICAM1;ADAMTSL1;COMP;ADAMTS4;ADAMTS5;ADAMTS2;CTSL;ADAMTS1;CTSK;SH3PXD2B;CAPN2;CYP1B1;COL10A1;QSOX1;TIMP1;CAPN1;ADAMTS7;POSTN;COL27A1;MMP2;P3H2;NPNT;HSPG2;DCN;MMP14;VCAN;COL2A1;LOX;COL6A2;ADAM12;COL6A1;COL8A1;COL6A3;ITGA5;GAS6;CD44;CD151;LAMA2;LAMA4;PAPLN;HTRA1;FURIN;ADAMTS12;THBS1;FBLN5;THSD4;SCUBE3;ACAN;HAS1;SPP1;TGFB1;FN1;COL1A1;COL1A2;SMOC1;ITGA10;ITGA11;MFAP2;COL9A3;TGFBI;RECK;FBN1* |
| External encapsulating structure organization (GO:0045229) | 59/217 | $2.83x10^{-24}$ | *VIT;ITGB1;COL18A1;ITGB5;LAMC3;ELN;SERPINE1;TNC;LAMC2;ICAM5;LAMC1;ICAM1;ADAMTSL1;COMP;TGM1;ADAMTS4;ADAMTS5;ADAMTS2;ADAMTS1;COL10A1;ADAMTS7;POSTN;COL27A1;MMP2;NPNT;HSPG2;DCN;MMP14;VCAN;COL2A1;LOX;COL6A2;ADAM12;COL6A1;COL8A1;COL6A3;ITGA5;CD44;LAMA2;LAMA4;PAPLN;FURIN;ADAMTS12;THBS1;FBLN5;THSD4;ACAN;SPP1;FN1;COL1A1;COL1A2;SMOC1;ITGA10;ITGA11;MFAP2;COL9A3;TGFBI;RECK;FBN1* |
| Extracellular structure organization (GO:0043062) | 58/216 | $1.08x10^{-23}$ | *VIT;ITGB1;COL18A1;ITGB5;LAMC3;ELN;SERPINE1;TNC;LAMC2;ICAM5;LAMC1;ICAM1;ADAMTSL1;COMP;ADAMTS4;ADAMTS5;ADAMTS2;ADAMTS1;COL10A1;ADAMTS7;POSTN;COL27A1;MMP2;NPNT;HSPG2;DCN;MMP14;VCAN;COL2A1;LOX;COL6A2;ADAM12;COL6A1;COL8A1;COL6A3;ITGA5;CD44;LAMA2;LAMA4;PAPLN;FURIN;ADAMTS12;THBS1;FBLN5;THSD4;ACAN;SPP1;FN1;COL1A1;COL1A2;SMOC1;ITGA10;ITGA11;MFAP2;COL9A3;TGFBI;RECK;FBN1* |

| | | | |
|---|---|---|---|
| skeletal system development (GO:0001501) | 36/158 | $1.06 \times 10^{-11}$ | *PKDCC;HHIP;NPR3;XYLT1;STC1;TNFRSF11B;PAPSS2;COMP;ADAMTS4;ACAN;GJA1;ANKH;TCOF1;FRZB;SH3PXD2B;COL10A1;CNMD;CMKLR1;TGM2;IGF2;SULF1;BMP6;SULF2;COL1A1;EXT1;RBP4;MMP14;VCAN;COL1A2;COL2A1;CDH11;CHI3L1;FGFR2;CD44;FBN1;FGFR1* |
| regulation of angiogenesis (GO:0045765) | 35/203 | $6.77 \times 10^{-08}$ | *ITGB1;FOXC1;RNH1;CXCL8;HSPB6;SERPINE1;TWIST1;HSPB1;GATA4;THBS2;THBS1;MTDH;NPPB;BMPER;ADAMTS1;CYP1B1;EMILIN1;CNMD;NIBAN2;PTGIS;SPHK1;ATP2B4;VEGFC;HMGA2;SULF1;KLF4;HSPG2;DCN;SFRP2;TNMD;ADAM12;FGF18;CTNNB1;CHI3L1;ANGPTL4* |
| positive regulation of angiogenesis (GO:0045766) | 23/116 | $5.18 \times 10^{-06}$ | *ITGB1;CXCL8;PTGIS;HSPB6;SPHK1;SERPINE1;TWIST1;HMGA2;HSPB1;VEGFC;GATA4;KLF4;THBS1;MTDH;SFRP2;BMPER;MDK;ADAM12;FGF18;CYP1B1;CHI3L1;ANGPTL4;ITGA5* |
| positive regulation of vasculature development (GO:1904018) | 19/102 | $1.49 \times 10^{-04}$ | *ITGB1;CXCL8;PTGIS;HSPB6;SPHK1;SERPINE1;TWIST1;HMGA2;HSPB1;VEGFC;GATA4;THBS1;MTDH;SFRP2;ADAM12;FGF18;CYP1B1;CHI3L1;ANGPTL4* |

**Table S5: Top five canonical pathways associated with DEGs identified in dimpled versus smooth samples at day 14 post-seeding generated using IPA**.

| Ingenuity canonical pathways | -log (*p*-value)* | z-score** | Ratio*** | Molecules |
|---|---|---|---|---|
| Regulation of Insulin-like Growth Factor (IGF) transport and uptake by IGFBPs | 11 | 3.21 | 14/124 | *APOB,APOE,CP,FN1,GAS6,IGF2,IGFBP2,IGFBP3,IGFBP4,IGFBP6,LTBP1,SPARCL1,TIMP1,TNC* |
| Osteoarthritis pathway | 11 | 3.21 | 18/238 | *ANKH,CCN4,CNMD,COL10A1,CXCL8,DCN,DKK1,FGF18,FN1,GLI1,HTRA1,IL1R1,ITGA11,ITGA5,ITGB5,PTCH1,RBP4,SPHK1* |
| Extracellular matrix organisation | 8.3 | 3.32 | 11/107 | *COL10A1,COL6A1,COL6A3,DCN,FN1,ITGA5,ITGB5,LAMA2,LAMC2,SERPINE1,TNC* |
| Post-translational protein phosphorylation | 8.3 | 2.71 | 11/107 | *APOB,APOE,CP,FN1,GAS6,IGFBP3,IGFBP4,LTBP1,SPARCL1,TIMP1,TNC* |
| Sheddase signalling pathway | 8.09 | 0.54 | 14/204 | *CCN3,FAS,FN1,HGF,IGF2,IGFBP2,IGFBP3,IGFBP4,IGFBP6,IT* |

| | | | | |
|---|---|---|---|---|
| | | | | *GA5,ITGB5,PLAUR,POSTN,TIMP1* |

*-log(*p*-value) reflects enrichment significance.
**z-score indicates the predicted activation or inhibition of each pathway.
***Ratio is an indicator of the proportion of pathway genes that overlap with the uploaded dataset, indicating its relevance.

*Abbreviations: APOB, Apolipoprotein B; APOE, Apolipoprotein E; CP, Ceruloplasmin; FN1, Fibronectin 1; GAS6, Growth Arrest-Specific 6; IGF2, Insulin-like Growth Factor 2; IGFBP2, Insulin-like Growth Factor Binding Protein 2; IGFBP3, Insulin-like Growth Factor Binding Protein 3; IGFBP4, Insulin-like Growth Factor Binding Protein 4; IGFBP6, Insulin-like Growth Factor Binding Protein 6; LTBP1, Latent Transforming Growth Factor Beta Binding Protein 1; SPARCL1, Secreted Protein Acidic and Rich in Cysteine-Like 1; TIMP1, Tissue Inhibitor of Metalloproteinases 1; TNC, Tenascin C; ANKH, Ankylosis Progressive Homolog; CCN4, Cellular Communication Network Factor 4; CNMD, Chondromodulin; COL10A1, Collagen Type X Alpha 1; CXCL8, C-X-C Motif Chemokine Ligand 8; DCN, Decorin; DKK1, Dickkopf WNT Signaling Pathway Inhibitor 1; FGF18, Fibroblast Growth Factor 18; GLI1, GLI Family Zinc Finger 1; HTRA1, High Temperature Requirement Factor A Serine Peptidase 1; IL1R1, Interleukin 1 Receptor Type 1; ITGA11, Integrin Subunit Alpha 11; ITGA5, Integrin Subunit Alpha 5; ITGB5, Integrin Subunit Beta 5; PTCH1, Patched 1; RBP4, Retinol Binding Protein 4; SPHK1, Sphingosine Kinase 1; COL6A1, Collagen Type VI Alpha 1; COL6A3, Collagen Type VI Alpha 3; LAMA2, Laminin Subunit Alpha 2; LAMC2, Laminin Subunit Gamma 2; SERPINE1, Serpin Family E Member 1; CCN3, Cellular Communication Network Factor 3; FAS, Fas Cell Surface Death Receptor; HGF, Hepatocyte Growth Factor; PLAUR, Plasminogen Activator, Urokinase Receptor; POSTN, Periostin.*

**Table S6: Thermal conditions used for cDNA synthesis and quantitative real-time PCR analysis.**

| Thermal conditions used in GS00482 Thermal Cycler (G-STORM) for cDNA synthesis | | | |
|---|---|---|---|
| **Cycling step** | **Temperature** | **Duration** | |
| Priming | 25°C | 5 minutes | |
| Reverse transcription | 42°C | 30 minutes | |
| Heat-inactivation | 85°C | 5 minutes | |
| Hold | 4°C | ∞ | |
| **Thermal conditions used for quantitative real-time PCR analysis** | | | |
| **Cycling step** | **Temperature** | **Duration** | **Cycles** |
| Enzyme activation | 95°C | 30 seconds | 1 |
| Denaturation | 95°C | 5 seconds | 35 |
| Annealing/extension | 64°C | 5 seconds | |
| Melting curve | 65-95°C (increment 0.5°C) | 5 seconds/step | 1 |

**Table S7: Fabrication parameters for generating GWL files for 2PP lithography.**

| | |
|---|---|
| **Slice parameters** | |
| Slicing distance | 0.8 |
| Simplification tolerance | 0.05 |
| Self-intersections | Fix |
| **Fill parameters** | |
| Hatching distance | 0.5 |
| Counter count | 0.0 |
| Base slice count | 2.0 |
| Hatching angle | Auto |
| **Parameters to create GWL file** | |
| **System initialization** | |
| Invert Z axis | 1.0 |
| **Writing configuration** | |
| Mode | Galvo scan Continuous |
| Piezo settling time | 10.0 |
| Galvo acceleration | 10.0 |
| Stage velocity | 200.0 |
| **Scan field offsets** | |
| X offset | 0.0 |
| Y offset | 0.0 |
| Z offset | 0.0 |
| **Writing parameters** | |
| Power scaling | 1.0 |
| **Solid hatch lines writing parameters** | |
| Laser power | 100 |
| Scan speed | 50000 |
| **Base writing parameters** | |
| Laser power | 100 |
| Scan speed | 50000 |
| Interface position | 0.5 |
